# *Pcdh20* is a POU2F3 target gene required for proper tuft cell microvillus organization

**DOI:** 10.1101/2025.03.18.644042

**Authors:** Katherine E. Ankenbauer, Yilin Yang, Chi-Yeh Chung, Leonardo R. Andrade, Sammy Weiser Novak, Brenda Jarvis, Wahida H. Ali Hanel, Jiayue Liu, Victoria Sarkisian, Neil Dani, Evan Krystofiak, Gaizun Hu, Seham Ebrahim, Bechara Kachar, Qizhi Gong, Geoffrey M. Wahl, Ken S. Lau, Jeffrey W. Brown, Uri Manor, Kathleen E. DelGiorno

**Affiliations:** Department of Cell and Developmental Biology, Vanderbilt University, Nashville, TN, USA; Epithelial Biology Center, Vanderbilt University Medical Center, Nashville TN, USA; Center for Computational Systems Biology, Vanderbilt University, Nashville TN, USA; Gene Expression Laboratory, The Salk Institute for Biological Studies, La Jolla, CA, USA; Laboratory of Genetics and Center for Chemical Biology and Proteomics, The Salk Institute for Biological Studies, La Jolla, CA, USA; Waitt Advanced Biophotonics Center, Salk Institute for Biological Studies, La Jolla, CA, USA; Biological and Physical Sciences Department, Columbus State Community College, Columbus, OH, USA; Center for Membrane and Cell Physiology, Department of Molecular Physiology and Biological Physics, University of Virginia School of Medicine, Charlottesville, VA, USA; Laboratory of Cell Structure and Dynamics, NIDCD, NIH, Bethesda, MD, USA; Department of Cell Biology and Human Anatomy, School of Medicine, University of California, Davis, CA, USA; Chemical and Physical Biology Program, Vanderbilt University, Nashville TN, USA; Department of Surgery, Vanderbilt University Medical Center, Nashville, TN, USA; Vanderbilt Ingram Cancer Center, Vanderbilt University Medical Center, Nashville, TN, USA; Vanderbilt Digestive Disease Research Center, Vanderbilt University Medical Center, Nashville, TN, USA; Division of Gastroenterology, Department of Medicine, Washington University in St. Louis, MO, USA; Department of Cell and Developmental Biology, University of California, San Diego, CA, USA

**Keywords:** ChIP-seq, hair cells, stereocilia, mechanotransduction

## Abstract

**BACKGROUND & AIMS:** Tuft cells play protective roles in infection, inflammation, and tumorigenesis through the secretion of cytokines and eicosanoids. Tuft cells are known for their tall, blunt microvilli, thought to be analogous to mechanosensory hair cell stereocilia; however, a functional role for the microvillar apparatus has not been identified. POU2F3 is the master regulator transcription factor for tuft cells, yet how POU2F3 drives formation of this unique structure is unknown. Here, we aimed to identify POU2F3 target genes and commonalities between tuft and hair cells to better understand this unique structure.

**METHODS:** POU2F3 ChIP-seq was performed on tuft cells and compared to the hair cell transcriptome. Tuft cell RNA-seq datasets were interrogated for hair cell structural and mechanosensory genes; expression was validated. Intestinal and gallbladder tuft cells were examined using multiple light and electron microscopy (EM) modalities. PCDH20 was knocked down in mouse models and ultrastructural analyses were performed. The tuft cell cytoskeleton was modeled using AlphaFold3 prediction.

**RESULTS:** Genes encoding structural and mechanosensory proteins common to both tuft and hair cells, including *Pcdh20*, were identified. Imaging localized PCDH20 to tuft cell microvilli and hair cell stereocilia. Genetic ablation of *Pcdh20* in mice resulted in structural defects in tuft cell microvilli, including loss of rigidity and organization. Molecular modeling suggests PCDH20 homodimers link adjacent microvilli.

**CONCLUSIONS:** *Pcdh20* is a POU2F3 target gene in tuft cells, critical to maintain the rigid microvillar apparatus. These findings, together with the shared expression of mechanosensory components like TMC1, support the hypothesis that tuft cells could have mechanosensory capabilities analogous to cochlear hair cells.

## INTRODUCTION

Tuft cells are rare chemosensory cells found throughout the respiratory and digestive tracts that are analogous to taste cells^1^. They are defined by their highly specific cytoskeletal structure of tall, blunt microvilli, thick, deep actin rootlets, and a well-developed tubulovesicular system that projects throughout the supranuclear cytoplasm^2^. Due to this unique structure, tuft cells have been identified by electron microscopy (EM) by many independent groups and received several names (e.g. fibrillovesicular, multivesicular, brush, caveolated cells) before more recent consensus settled on ‘tuft cell’^2, 3^. Despite characterization of this cell type as early as 1956^4^, little functional data existed in the gastrointestinal tract until 2016 when three independent groups demonstrated that tuft cells function in the clearance of Helminth parasites in the intestines through the synthesis and secretion of cytokine IL-25^5-7^. Since then, tuft cells have been shown to sense the metabolite succinate in the intestine and trachea^8^ and to secrete cysteinyl leukotrienes in response to *Alternaria* challenge in the tracheal and airway epithelium^9^. In the gallbladder, tuft cells also produce cysteinyl leukotrienes as well as acetylcholine, driving contraction and release of mucins, which may play a role in controlling tissue inflammation in response to commensal microbes^10, 11^. In the pancreas, we have shown that tuft cells mitigate inflammation and inhibit tumorigenesis through eicosanoid prostaglandin D_2_^12^. Despite these varied and context-specific functions, tuft cells in different organs and disease states all require transcription factor POU2F3 and share a well-preserved structure, for which a functional role has yet to be demonstrated^13^.

POU Class 2 Homeobox 3 (POU2F3, also known as SKN-1A or OCT11) is the master regulator transcription factor for tuft cell formation^14^. Full body POU2F3 knockout mice lack tuft cells in all organs as well as electrophysiological and behavioral responses to sweet, umami, and bitter compounds, due to a complete loss of sweet, umami, and bitter taste cells^15^. Like taste cells, tuft cells express the taste chemosensation pathway including G protein coupled taste receptors^16^, which signal through GNAT3, PLCβ2, and ITPR3 to induce calcium flux, opening cation channel TRPM5; in taste cells this results in depolarization and secretion of ATP to adjacent nerves^17^. Outside of the tastebud, multiple studies have demonstrated that this machinery allows tuft cells to monitor intraluminal homeostasis, to achieve local responses to irritants, infection, or tumorigenesis via secreted effectors^1^. Unlike taste cells, tuft cells possess a distinct microvillus apparatus, currently without a known function.

Ultrastructural studies of murine gallbladder tuft cells show that, at a diameter of 0.18-0.2 μm, tuft cell microvilli are thicker than the conventional microvilli characteristic of intestinal enterocytes. Due to their stiff appearance, tuft cell microvilli, instead, more closely resemble the stereocilia of hair cells although not exhibiting the staircase morphology^3^. Freeze fracture deep-etching replicas and EM studies are consistent with this assessment and led to the hypothesis that, like the stereocilia of hair cells, tuft cell microvilli could also function in mechanotransduction^3^.

In the inner ear, auditory and vestibular hair cells detect sound and head position by converting mechanical stimuli into electrical signals through actin-rich stereocilia organized into bundles. These bundles are held together by tip-link filaments composed of cadherin 23 (CDH23) and protocadherin 15 (PCDH15), which coordinate bundle architecture and enable mechanotransduction^18^. Tension on the stereocilia links from mechanical cues opens a channel composed of transmembrane channel-like proteins 1 and 2 (TMC1, TMC2), located on the tips of shorter row stereocilia^19^. The resulting influx of Ca^2+^ and K^+^ ions depolarizes the hair cells and results in the secretion of neurotransmitters to afferent nerve fibers^20^. Disruption of stereocilia structure or this carefully controlled mechanotransduction signaling process often results in deafness or balance disorders; highlighting a critical way in which form follows function^19^.

In the intestine, similar adhesion systems are formed by trans-heterophilic interactions between the protocadherins CDHR2 and CDHR5, which assemble intermicrovillar adhesion complexes (IMACs)^21^. Tuft cells and enterocytes express these IMACs, which are critical for organizing microvilli within the brush border^22^. IMACs can form within single cells or can form links between neighboring cells, which promotes proper spacing and packing of the microvilli^23-25^. Loss of CDHR2 and CDHR5 leads to brush border defects such as microvillus shortening^26, 27^. Ablation of CDHR2 in mouse models leads to disorganized microvillus packing^26^, whereas loss of CDHR5 produces more pronounced brush border defects and severe mucosal damage in DSS-induced colitis^27^. Taken together, these findings in both the inner ear and the intestine highlight that adhesion complexes do not simply organize actin-rich protrusions but are essential for enabling the specialized functions these epithelial structures support.

The structural similarities between tuft cell microvilli and hair cell stereocilia led us to ask if tuft cells express components of the mechanotransduction machinery and how POU2F3 regulates this process. Here, we performed POU2F3 chromatin immunoprecipitation sequencing (ChIP-seq) on isolated tuft cells and identified structural gene targets of POU2F3 that are shared with hair cells. Protocadherin 20 (PCDH20) was identified as a novel structural protein in both cell types and a likely component of intermicrovillar linkage in tuft cells. Genetic ablation of PCDH20 resulted in impaired microvillar organization and fewer secretory vesicles in tuft cells. Collectively, we have identified a critical tuft cell structural protein as a first step to understanding how this unique tuft cell structure dictates cell function.

## RESULTS

### ChIP-seq identifies POU2F3 target genes in tuft cells

Transcription factor POU2F3 is the master regulator of tuft cell formation in normal organs, as well as for those tuft cells that arise *de novo* in disease, such as in pancreatic tumorigenesis^5, 12, 28, 29^. POU2F3 is known to function in taste sensation and keratinocyte maturation^15, 30^, however little is known about how POU2F3 regulates tuft cell formation. To identify targets of POU2F3 binding, we performed chromatin immunoprecipitation sequencing (ChIP-seq) on isolated small intestinal tuft cells. Tuft cells make up only ∼0.4% of the epithelium in the murine small intestines^5^. Therefore, to acquire the requisite 10,000 cells needed for this analysis, we sorted large portions of normal duodenum from multiple adult mice and pooled samples. Tuft cells were isolated and concentrated by fluorescence activated cell sorting (FACS) using a Siglec F^+^;EpCAM^+^ signature, as previously described^5, 12, 31^. Duplicate POU2F3 ChIP-Seq libraries were prepared from two independent experiments and compared to an untargeted IgG binding control (**Figure 1A**). Q enrichment score (QES) analysis was performed to verify the quality of these POU2F3 binding sites. Only peaks (FDR < 0.05) common between the two biological replicates were called and any that overlapped with the IgG control and/or ENCODE blacklist regions were removed. Compared with the reference values for the quality metrics generated from 392 data sets from ENCODE, the QES from our POU2F3 ChIP-seq was ranked at the level of moderate high to very high, suggesting high quality POU2F3 peaks.

**Figure 1.**
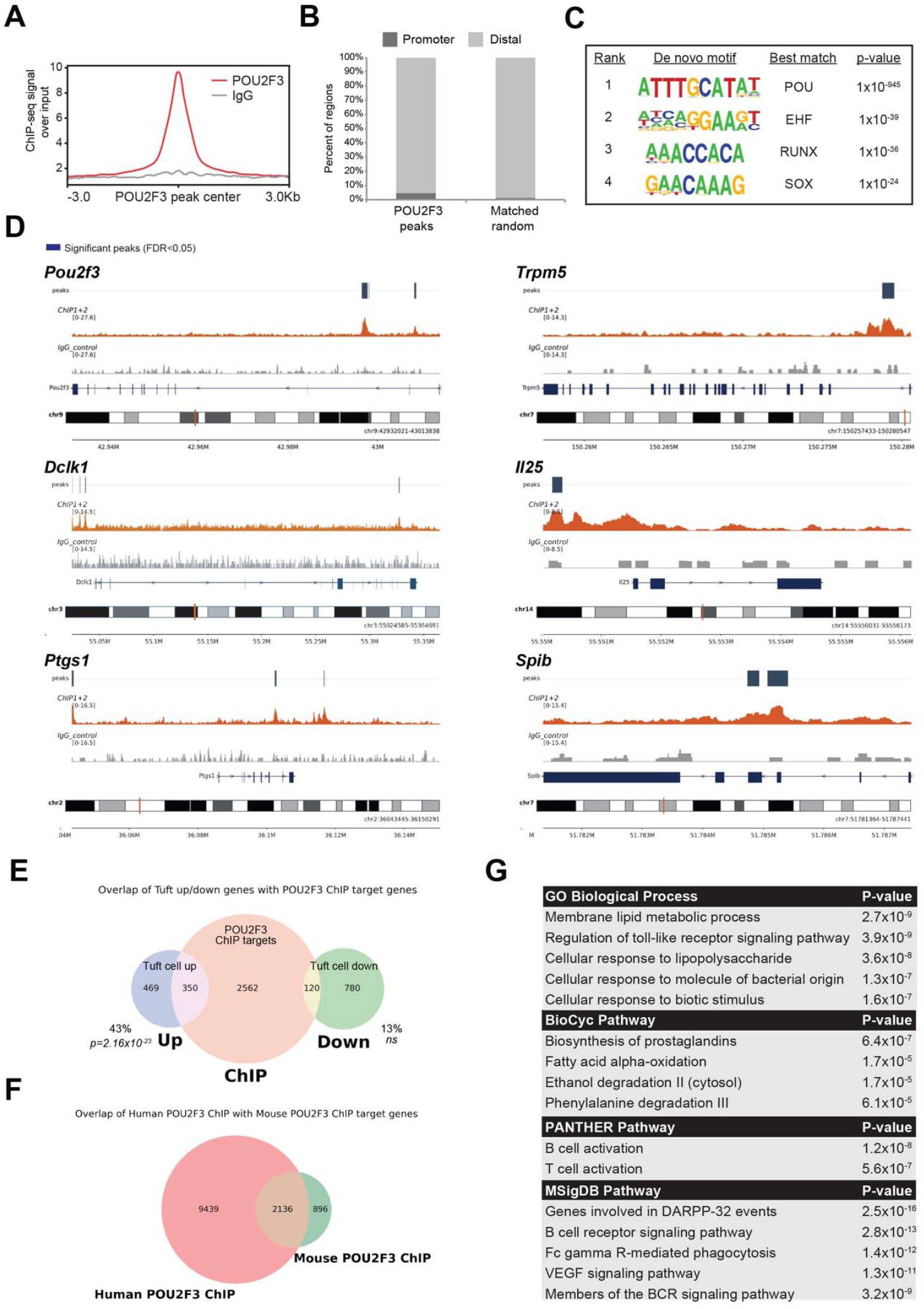
ChIP-seq identifies POU2F3 target genes in tuft cells. **(A)** POU2F3 and IgG control ChIP-seq signal. n = 2 replicates of multiple pooled male mice. (**B**) Bar graph demonstrating enrichment of POU2F3 peaks in gene promoters. (**C**) Motif analysis identifying likely POU2F3 consensus binding sequences (**D**) Tracks showing POU2F3 binding sites in the genes of canonical tuft cell markers. (**E**) Venn diagram of POU2F3 ChIP-seq target genes and differentially expressed genes identified as either up- or down-regulated in duodenal tuft cells as compared to non-tuft epithelial cells by RNA-seq^12^. (**F**) Venn diagram of POU2F3 ChIP-seq target genes in either normal murine tuft cells or in a human cell line of a tuft-like variant of non-small cell lung cancer^34^.(**G**) Genomic Regions Enrichment of Annotations (GREAT) analysis of POU2F3 ChIP-seq target genes.

Next, the distribution of peaks across functional domains in the genome was analyzed. POU2F3 peaks were strongly enriched in gene-associated functional domains, such as promoter (defined as -1 kb to +100 bp of transcription start site, 14.27%), un-translated region (2.45%), intron (36.31%), and exon (4.43%), where they usually mapped within 1kb of the transcription start site (TSS), indicating that the peaks generated from our POU2F3 ChIP-seq were not located randomly on the genome but instead were associated with core promoters (**Figure 1B, S1A**). To identify direct POU2F3 transcriptional targets, we defined genes that have POU2F3 binding sites as proximal, 5 kb upstream or 1kb downstream of the TSS, or distal, up to 1000 kb, as POU2F3-associated genes. We identified a total of 2828 POU2F3 high confidence binding regions which were mapped to 3032 genes across all 21 genomic chromosomes (**Figure S1B**).

Next, we performed *de novo* motif analysis for our POU2F3 ChIP-seq data using HOMER. As expected, POU2F3 bound chromatin was significantly enriched for the consensus POU motif (p < 1 x 10^-945^), demonstrating that direct binding sites for POU transcription factors are enriched in our ChIP-seq data (**Figure 1C**). Interestingly, the consensus DNA binding motifs of several transcription factors highly expressed in tuft cells matched binding sequences derived *de novo* from POU2F3 ChIP-seq analysis. Included are *Ehf* (EHF, p < 1 x 10^-39^)^32^, *Runx1* (RUNX, p < 1 x 10^-36^), and *Sox9* (SOX, p < 1 x 10^-24^)^33^, suggesting the possibility that these factors bind cooperatively with POU2F3 to regulate tuft cell formation (**Figure 1C**)^12^.

To identify direct POU2F3 transcriptional targets in tuft cells, we compared our new POU2F3 ChIP-seq analysis with a previous messenger RNA (mRNA)-based dataset we generated using SmartSeq2 on low abundance samples where we identified 819 significantly upregulated and 900 significantly downregulated genes in tuft cells as compared to non-tuft epithelial cells in the duodenum^12^. When comparing the two datasets, we identified significant overlap of POU2F3 ChIP-seq target genes and the tuft cell transcriptomic signature (43%, p = 2.16 x 10^-23^), consistent with a role for POU2F3 in specifying tuft cell fate. Many POU2F3 target genes serve as canonical tuft cell markers, including *Trpm5, Dclk1, Ptgs1, Il25, Spib*, and *Pou2f3* itself (**Figure 1D, File S1**). Additional genes common to both datasets include *Alox5, Bmp2, Cd300lf, Chat, Gnat3, Hpgds, Sucnr1, Vav1*, and many others previously described to be enriched in tuft cells (**Figure 1E, File S2**)^12^. POU2F3 expression has also been described in a variant of human non-small cell lung cancer (NSCLC) expressing tuft cell markers^34^. To compare POU2F3 target genes between species and disease states, we compared our POU2F3 ChIP-seq dataset to a ChIP-seq dataset generated from a tuft cell-like human NSCLC cell line^34^. This comparison allows for the identification of a core, conserved POU2F3-driven transcriptional program independent of species or tissue context (normal vs. malignant). Interestingly, we found that 70.4% (2136/3032) of POU2F3 ChIP target genes identified in murine tuft cells overlapped with the human dataset, which also included *ALOX5, BMP2, DCLK1, GNAT3, HPGDS, SUCNR1*, and *VAV1*, consistent with a role for POU2F3 in driving a tuft-like fate (**Figure 1F, File S3**).

To gain broader insight into the role of POU2F3 target genes, Genomic Regions Enrichment of Annotations (GREAT) analysis was performed on our POU2F3 ChIP-seq dataset. Here we identified immune cell signaling pathways such as ‘B cell receptor signaling pathway’, ‘B cell activation’, ‘Members of the BCR signaling pathway’, and ‘T cell activation’. Lipid metabolism pathways, including ‘Fatty acid alpha-oxidation’ and ‘Biosynthesis of prostaglandins’, were identified, consistent with a known role for tuft cells in eicosanoid synthesis^12^. Finally, we identified several pathways consistent with a possible role for tuft cells in microbial interactions, including ‘Cellular response to lipopolysaccharide’, ‘Cellular response to molecule of bacterial origin’, and ‘Cellular response to biotic stimulus’ (**Figure 1G**). Gene Ontology (GO) pathway analysis of genes common to both our ChIP-seq and RNA-seq datasets again identified immune cell signaling pathways such as ‘B cell receptor signaling pathway’ and ‘Regulation of B cell receptor signaling pathway’ and eicosanoid biosynthesis pathways such as ‘Synthesis of lipoxins’ (**Figure S2**). Collectively, these analyses identified genes in tuft cells putatively regulated by POU2F3. Significant overlap between ChIP target genes and mRNA enriched in tuft cells identifies direct POU2F3 target genes and a mechanism by which POU2F3 drives tuft cell fate.

### POU2F3 ChIP-seq reveals a cochlear hair cell gene signature

Tuft cells are identified by their distinct cytoskeletal structure of tall, blunt microvilli and deep actin rootlets (**Figure 2A**). Among POU2F3 target genes in our ChIP-seq analysis, we identified several encoding known tuft cell structural proteins including *Avil*^*35*^, *Cdhr2 and Cdhr5*^*22*^, and *Krt18* and *Vil1*^*36*^ (**File S1**). These data suggest that POU2F3 activity is required for assembly of the microvillus/rootlet apparatus. Freeze fracture electron microscopy studies of murine gallbladder tuft cells suggest that tuft cell microvilli are more similar to cochlear hair cell stereocilia than the conventional microvilli characteristic of intestinal enterocytes^3^. Based on size and ultrastructural features suggesting stiffness, these data indicate a possible role for tuft cell microvilli in mechanotransduction. To identify commonalities between tuft cells and cochlear hair cells, a prototypical mechanosensory cell type, we compared our POU2F3 ChIP-seq dataset to a single cell RNA-seq (scRNA-seq) dataset generated from adult murine cochlea^37^. As shown in **Figure 2B**, we found that 27.2% (825/3032) of ChIP-seq target genes overlapped with the inner hair cell transcriptomic signature. Mutations in many of these genes, including *Grxcr1*^*38*^, *Cldn14*^*39*^, and *Myo6*^*40*^ are associated with deafness, highlighting their critical role in hair cell function (**Figure 2C, S3**).

**Figure 2.**
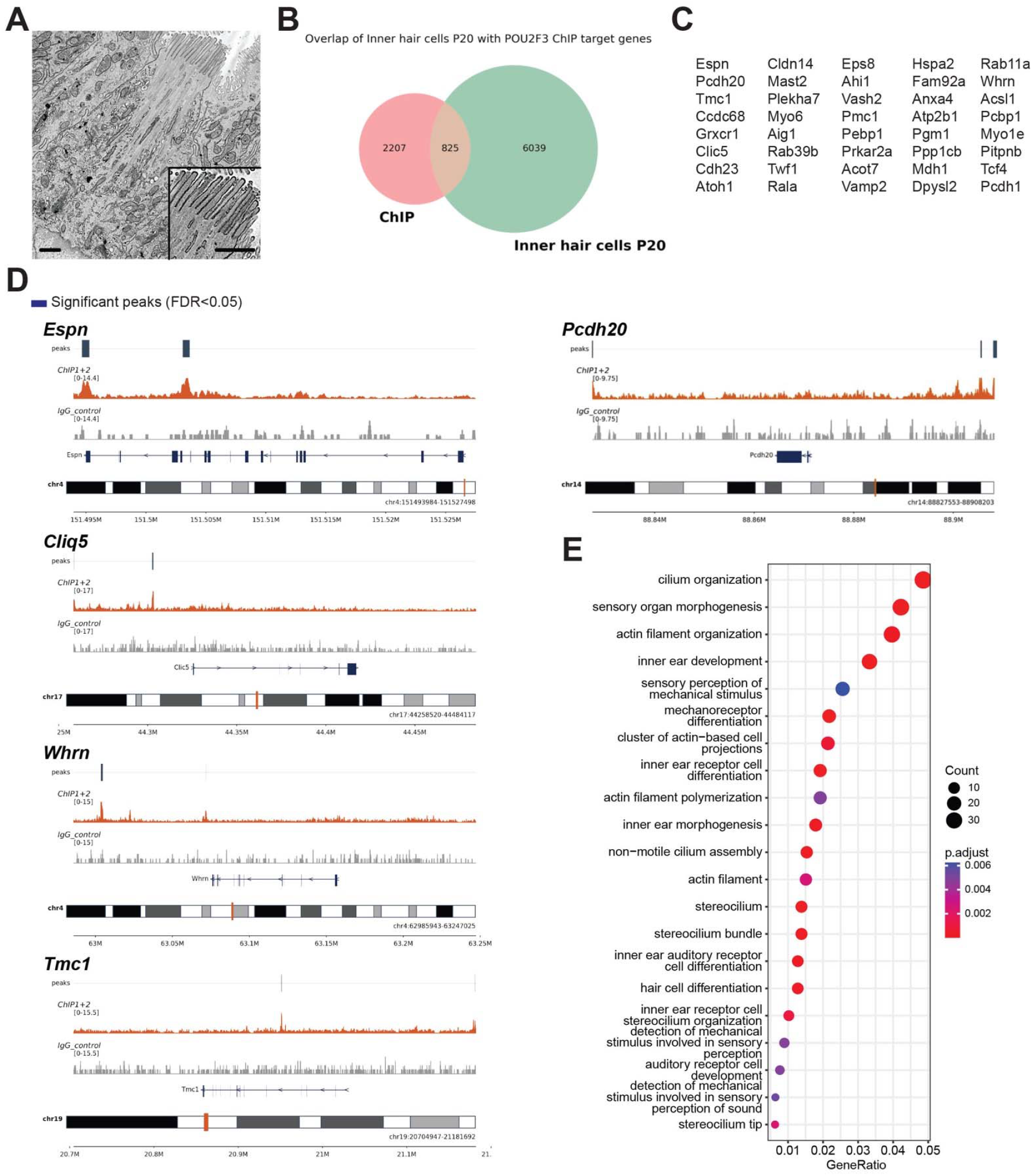
POU2F3 ChIP-seq reveals a cochlear hair cell transcriptomic signature. (**A**) Scanning electron microscopy of a thin tissue section of a small intestinal tuft cell, highlighting the microvillar structure. Scale bars, 1 μm. (**B**) Venn diagram of POU2F3 ChIP-seq target genes and a murine cochlear hair cell RNA-seq dataset generated in a study by Jean et al^37^. (**C**) Select genes from the 825 common to both datasets in (B). (**D**) Tracks showing POU2F3 binding sites in tuft cells for genes expressed in cochlear hair cells including known markers (*Espn, Clic5, Whrn, Tmc1*) and protocadherin 20 (*Pcdh20*). (**E**) Gene Ontology (GO) analysis of the 825 genes common to both the intestinal tuft cell POU2F3 ChIP-seq dataset and the hair cell RNA-seq dataset, highlighting structure-function pathways.

Among genes common to both our tuft cell POU2F3 ChIP-seq dataset and the outer hair cell transcriptome, we identified *Espn*, which is an actin cytoskeletal regulatory protein that contributes to the integrity of parallel actin bundles in the microvilli and stereocilia of chemosensory and mechanosensory cells^41, 42^. *Clic5*, a member of the chloride intracellular channel protein family, is required for stereocilia formation in the inner ear and normal hearing^43-45^. The gene *Whrn*, encoding Whirlin, is necessary for elongation and maintenance of inner and outer hair cell stereocilia in the organ of Corti and defects are also associated with deafness^46^. *Tmc1* is a pore-forming member of the mechanotransducer (MET) non-selective cation channel complex located at the tips of stereocilia that mediates sensory transduction. Loss or mutation also results in deafness by disturbing induction of the tension-induced MET channel and downstream signaling^47^. Finally, we identified *Pcdh20*, or protocadherin 20; expression of which has not been previously described in either tuft or hair cells (**Figure 2D**). Protocadherins are integral membrane proteins that mediate calcium-dependent cell-cell adhesion and are important components of the MET complex in the stereocilia of hair cells^48, 49^. PCDH20 is a type-I single-pass transmembrane protein belonging to the delta-2 protocadherin subgroup, containing seven extracellular cadherin (EC) domains, though its capacity for cell-cell adhesion has not been directly demonstrated^50, 51^. While functional data on the role of PCDH20 in cochlear hair cells is lacking, Genome-Wide Association Analysis (GWAS) on normal hearing function (hearing thresholds and pure-tone averages) in populations in Italy, Central Asia, the United Kingdom, and Finland identified significant loci close to PCDH20 as candidates for progressive hearing function and loss^52^. Interestingly, we also identified *ESPN, CLIC5, WHRN, TMC1*, and *PCDH20* as POU2F3 gene targets in the ChIP-seq study performed on the tuft cell-like human NSCLC cell line (**File S3**)^34^.

To gain deeper insight into features shared between tuft and hair cells, and to identify a possible role for the tuft cell microvilli, we performed GO analysis on the 825 genes shared between our POU2F3 ChIP-seq dataset and the adult inner hair cell transcriptome. Cytoskeletal pathways identified included ‘cilium organization’, ‘actin filament organization’, ‘actin filament polarization’, and ‘stereocilium’. Similarities in cell type formation include ‘inner ear development’, ‘inner ear morphogenesis’, and ‘inner ear auditory receptor cell differentiation’. Finally, similarities in function between tuft cells and hair cells are suggested by the terms ‘sensory perception of mechanical stimulus’, ‘stimulus involved in sensory perception’, and ‘detection of mechanical stimulus involved in sensory perception of sound’ (**Figure 2E, File S4**). Collectively, these data suggest that POU2F3 in tuft cells controls transcription of genes known to be involved in stereocilia formation and function.

### PCDH20 is a tuft cell-specific protocadherin in the gut epithelium

To evaluate broad expression of identified cochlear hair cell genes in diverse tuft cell populations, we first analyzed several publicly available RNA-seq datasets. In a study conducted by Nadjsombati et al., normal tuft cell populations derived from murine small intestines, colon, gallbladder, thymus, and trachea were concentrated by FACS using an IL-25 fluorescent reporter mouse model and underwent bulk RNA-sequencing^8^. As shown in the principal component analysis (PCA) in **Figure 3A**, tuft cell populations clustered by tissue type. *Espn* and *Clic5* expression were found to be significantly elevated in tuft cells as compared to non-tuft epithelial populations in the small intestines and were highly expressed in all tuft cell populations from all organs assayed. *Pcdh20* expression was absent from non-tuft intestinal cells but was identified at varying levels of expression in all tuft cell populations (**Figure 3B**). *Whrn* was expressed at very low levels and *Tmc1* was not identified.

**Figure 3.**
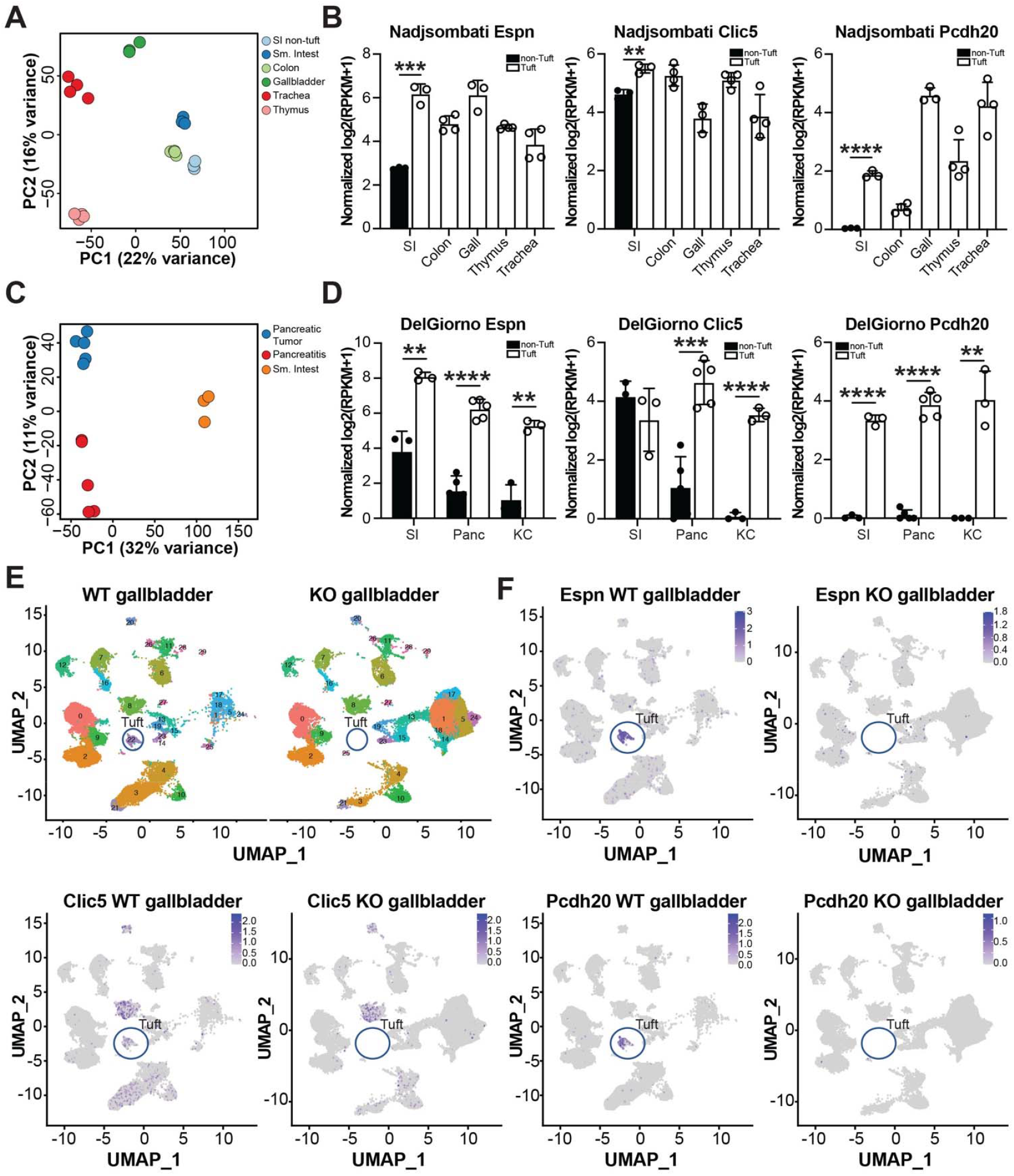
PCDH20 is a tuft cell-specific protocadherin in the gut epithelium. (**A**) Principal component analysis (PCA) comparing the transcriptomes of tuft cells from the murine small intestines, colon, gallbladder, thymus, and trachea, and non-tuft intestinal epithelial cells generated by the study from Nadjsombati et al (n = 3-4 mice/condition)^8^. (**B**) Expression of *Espn, Clic5*, and *Pcdh20* in the datasets in (A). (**C**) PCA comparing the transcriptomes of tuft cells from the normal murine duodenum, caerulein-induced chronic pancreatitis, or *Kras*^*G12D*^-induced pancreatic neoplasia generated by studies from DelGiorno et al (n = 3-5 mice/condition)^12, 31^. (**D**) Expression of *Espn, Clic5*, and *Pcdh20* in the datasets in (C). (**E**) Uniform manifold approximation and projection (UMAP) of murine gallbladder scRNA-seq collected from either *Pou2f3+/+* or *Pou2f3*-/- mice generated by a study from O’Leary et al^11^. (**F**) UMAPs of *Espn, Clic5*, or *Pcdh20* expression in either *Pou2f3+/+* or *Pou2f3*-/- gallbladders. Data are represented as mean +/-SD. ^**^, p < 0.01; ^***^, p < 0.005; ^****^, p < 0.001.

Next, we analyzed bulk RNA-seq datasets generated from metaplasia-derived, disease associated tuft cells. Here, DelGiorno et al., isolated tuft cells using the FACS strategy of collecting Siglec F^+^ EpCAM^+^ populations from the pancreata of mice with either chronic pancreatitis or oncogenic *Kras*^*G12D*^-induced neoplasia^12, 31^. As shown in **Figure 3C**, tuft cell populations clustered by disease state, however pancreas-derived tuft cells clustered more closely to each other than to normal duodenal tuft cells. In these datasets, *Espn* expression was significantly higher in tuft cells, as compared to non-tuft epithelial cells in both disease states, consistent with expression patterns identified in **Figure 3B**. *Clic5* expression was found to be significantly enriched in pancreas-derived tuft cells as compared to non-tuft cells; expression was present in this duodenal tuft cell dataset but was not significantly elevated. Finally, we assayed for *Pcdh20* expression and identified tuft cell-specific expression in all datasets (**Figure 3D**). Again, *Whrn* and *Tmc1* were not detected.

Finally, we evaluated expression of cochlear hair cell genes in several scRNA-seq datasets. In O’Leary et al., the authors performed scRNA-seq on gallbladders derived from wild type *Pou2f3+/+* control mice or from *Pou2f3*-/- mice, which cannot form tuft cells (**Figure 3E**)^5, 28^. As shown in Figure 3F, expression of *Espn, Clic5*, and *Pcdh20* were all found in tuft cells with *Espn* and *Pcdh20* expression largely lost in *Pou2f3*-/- mice. Again, *Whrn* and *Tmc1* expression was not identified (**Figure S4A**). Similar results were obtained in a scRNA-seq dataset of murine pancreatitis tuft cells (**Figure S4B**). Altogether, these data identify expression of several cochlear hair cell genes at the mRNA level in tuft cells with *Pcdh20* being largely tuft cell specific in the gut epithelium.

### PCDH20 is enriched in tuft cell microvilli

To validate expression of cochlear hair cell markers at the protein level in tuft cells, we performed immunostaining in both the murine gallbladder and small intestines. Tuft cells were identified in both tissues by F-actin staining which labels the tuft cell microvilli and deep actin rootlets characteristic of this cell type, as in previous studies^31, 53^. Consistent with previous work in the small intestines, Espin (*Espn*) was identified in tuft cells^54^. We show here that Espin is expressed in gallbladder tuft cells as well, and that expression largely overlaps with F-actin, consistent with a role in actin bundling (**Figure 4A-B**)^55^. Next, we evaluated PCDH20 expression in tuft cells by immunostaining. To validate PCDH20 antibody specificity, we generated controls by transfecting U2OS and HeLa cells with either a GFP or a PCDH20-GFP construct and performed immunostaining for PCDH20 (**Figure S5A-H**). Once PCDH20 expression was confirmed only in PCDH20-GFP transfected cells, we performed immunofluorescence on duodenal, gallbladder, and *Kras*^*G12D*^-induced neoplastic pancreas tuft cells. In all tuft cell populations, PCDH20 localized to the apical microvilli, indicating a possible role in intermicrovillar linkage (**Figure 4C-D, S5I-L**). Given the identification of *PCDH20* as a candidate hearing loss gene and its shared expression between tuft and hair cells, we next confirmed its localization in the inner ear. We performed immunofluorescence for PCDH20 in the cochlear inner hair cells of adult wild-type mice. Consistent with a role as an interstereociliary link protein, PCDH20 localized to the stereocilia of the inner hair cells (Figure S6A). These data validate that PCDH20 is a shared structural component of both tuft cell microvilli and hair cell stereocilia.

**Figure 4.**
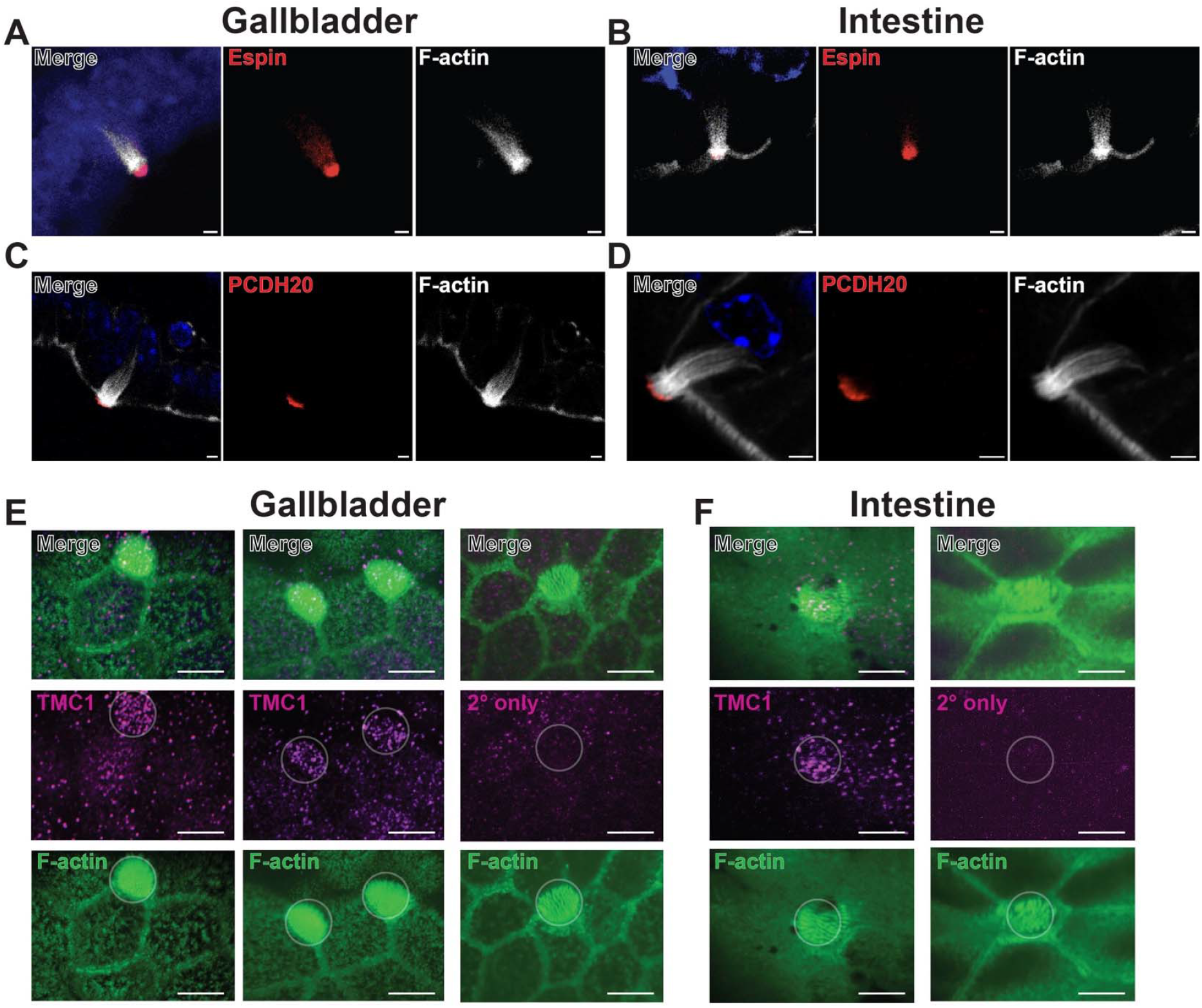
PCDH20 localizes to tuft cell microvilli. Immunofluorescence (IF) staining of tuft cells in the murine small intestines or gallbladder for F-actin (phalloidin, white), DAPI (blue) and either (**A-B**) Espin or (**C-D**) PCDH20, scale bars 2 μm. IF for TMC1 (magenta) and F-actin (phalloidin, green) in tuft cells in the (**E**) gallbladder or (**F**) small intestines. Scale bars, 5 μm. 2° only, secondary antibody staining only. Representative images reflect analyses performed on a minimum of n = 3 mice.

Finally, we examined tuft cells for expression of Whirlin (*Whrn*) and TMC1, which were identified in our ChIP-seq analysis, but were not detected in our analyses of published RNA-seq datasets. Whirlin was identified in gallbladder, but not intestinal tuft cells, with expression enriched in the microvilli (**Figure S7**). Interestingly, expression of TMC1, a member of the MET complex that mediates sensory transduction in the hair cells, was identified in the microvilli of both gallbladder and intestinal tuft cells (**Figure 4E-F**). Collectively, these analyses confirm expression of cochlear hair cell markers in tuft cells, with several localized to the microvillar apparatus.

### Tuft cells are characterized by extensive intermicrovillar linkages

In cochlear hair cells, stereocilia are connected by tip links, or small filaments required for the transmission of mechanical signals and hearing. These tip links are constructed of two proteins, cadherin 23 and protocadherin 15 (PCDH15). Loss of either result in deafness in murine models^56, 57^. Tuft cells have been shown to express CDHR2 (PCDH24) and 5, leading us to ask whether PCDH20 could play an analogous role in tuft cells^22^.

To determine whether tuft cell microvilli are interconnected by protein links, we applied several orthogonal EM techniques to resolve these structures in both the murine intestines and gallbladder. In the gallbladder, we performed both scanning electron microscopy (SEM) and transmission electron microscopy (TEM). We identified intermicrovillar links by SEM (**Figure 5A-C**), as well as TEM (**Figure 5D**), which also revealed links between tuft cell microvilli and neighboring epithelial cells (**Figure 5E**). In the small intestines, we found that the glycocalyx made these structures difficult to resolve by SEM (**Figure 5F-H**). To ameliorate this issue, we took a higher resolution approach and performed freeze fracture deep etching electron microscopy (FFE-EM), which led to the discovery that the tuft cell microvillar apparatus is characterized by extensive linkages between individual microvilli, consistent with a predicted rigidity attributed to the structure (**Figure 5I-J**). While the luminal surface is often coated in mucus, the fine, uniform, thread-like nature of these connections, especially as resolved by high-resolution freeze fracture EM (**Figure 5I-J**), is consistent with proteinaceous intermicrovillar links^58^. Collectively, these analyses reveal intermicrovillar linkages in tuft cells, which may limit microvillar deflection or mobility—similar to the constraints seen in hair cells—and could support a mechanotransduction role by stabilizing a MET-like complex.

**Figure 5.**
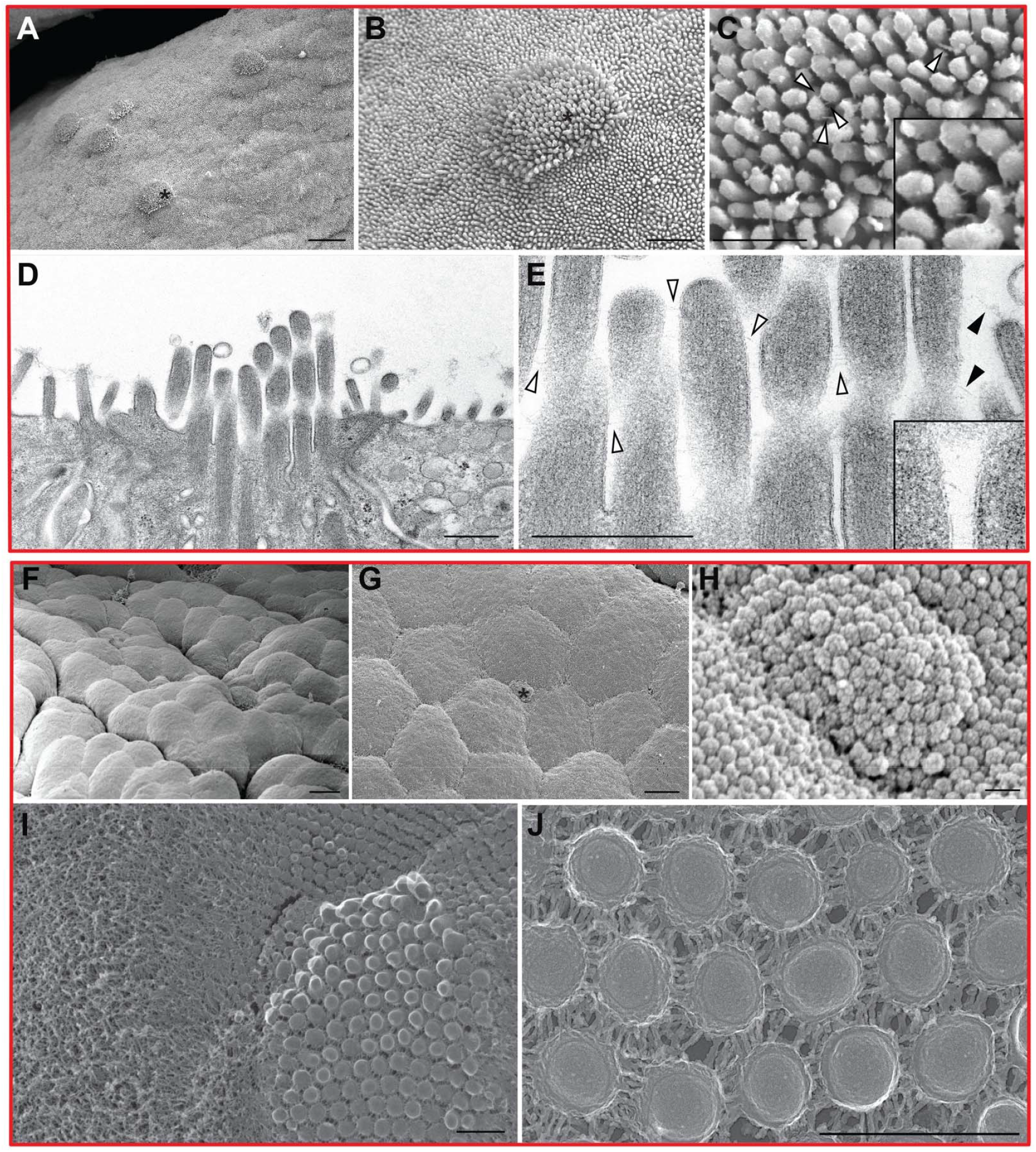
High resolution ultramicroscopy reveals intermicrovillar linkages in tuft cells. (**A**) SEM of the apical surface of the murine gallbladder, scale bar, 6 μm and (**B-C**) a tuft cell highlighting links between microvilli. Scale bars, 2 μm and 1 μm, respectively. (**D-E**) Transmission electron microscopy (TEM) of a gallbladder tuft cell highlighting links between tuft cell microvilli as well as between the tuft and a surrounding cell. Scale bars, 400 nm. White arrows, intermicrovillar tuft cell links. Black arrows, links between tuft and neighboring cell. Scanning electron microscopy (SEM) of (**F**) a normal intestinal villus at low magnification, scale bar, 3 μm, and (**G-H**) a tuft cell highlighting the apical microvilli, scale bars, 2 nm. (**I-J**) Freeze etch electron microscopy revealing extensive intermicrovillar linkages in intestinal tuft cells. Scale bars, 500 nm. SEM reflects analyses performed on a minimum of n = 3 mice. TEM and Freeze etch, n = 1 mouse.

### PCDH20 loss impairs proper tuft cell microvillar organization

By immunostaining, we identified PCDH20 expression in tuft cell microvilli, and our EM analyses revealed extensive intermicrovillar linkages in intestinal and gallbladder tuft cells (**Figures 4-5**). To determine if PCDH20 localizes to microvilli in a pattern suggestive of lateral linkage, we performed EM immunogold staining. As shown in **Figure 6A**, PCDH20 localized to tuft microvilli, consistent with immunostaining. To determine if PCDH20 plays a functional role in the formation of the tuft cell microvillus, we generated a genetically engineered mouse model (GEMM) of PCDH20 loss using clustered regularly interspaced short palindromic repeats (CRISPR). Multiple founder mice were generated and screened for PCDH20 loss by DNA sequencing (**Figure S8**) and (real time) polymerase chain reaction (**Figure S9A-B**). Ultimately, we chose to further assay a mouse with a 121 base pair deletion and an 11 base pair insertion in exon 3 of *Pcdh20* (**Figure S8**). Loss of PCDH20 expression in *Pcdh20-/-* mice was confirmed by immunostaining hair cells and intestinal tuft cells (**Figure 6B, S6B, S9C**). Ultrastructural analyses were further pursued in the gallbladder due to the high density of tuft cells in this organ as compared to the intestines.

**Figure 6.**
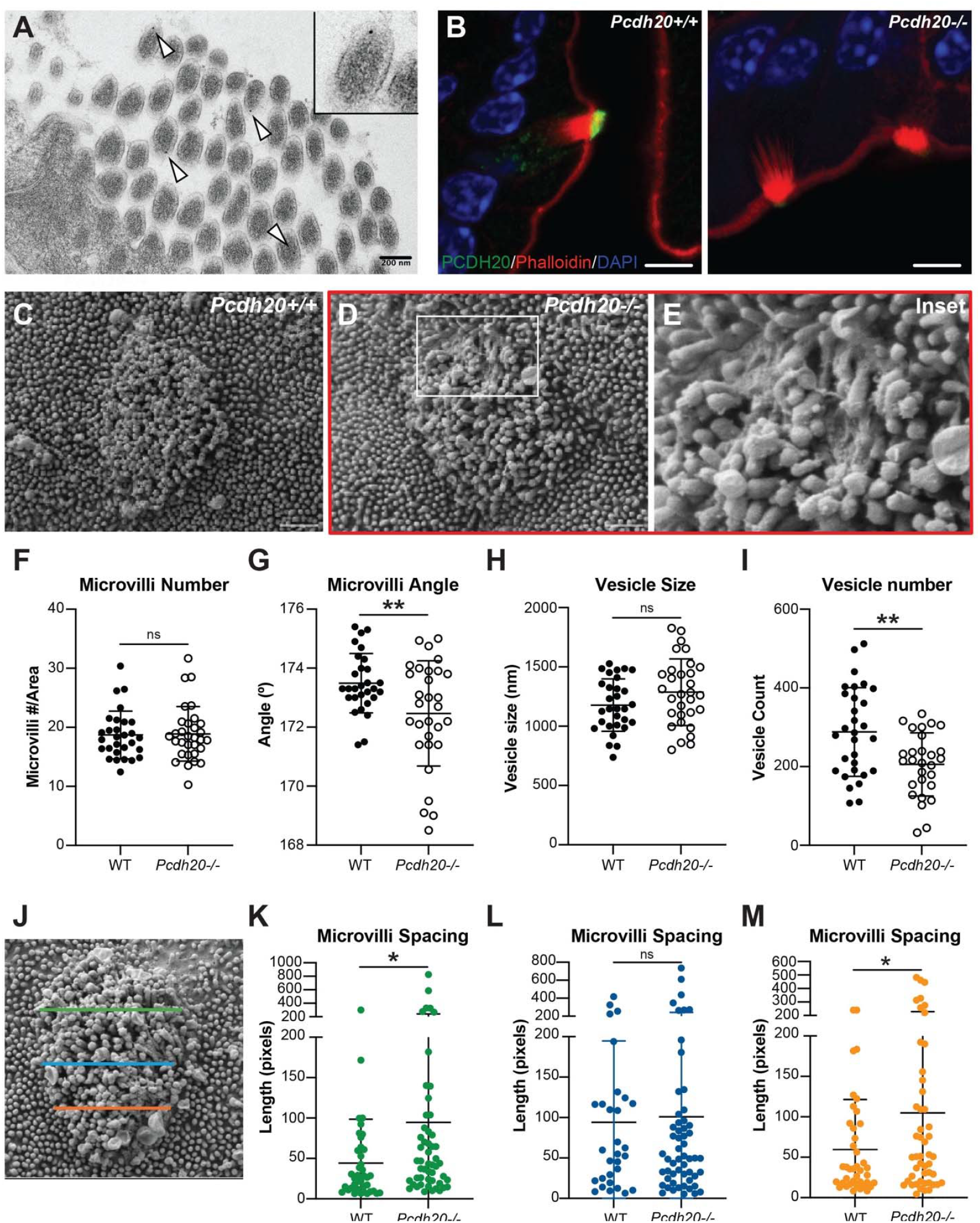
PCDH20 maintains tuft cell microvillar structure and rigidity. (**A**) Immunogold labeling of PCDH20 in a gallbladder tuft cell. Scale bar, 200 nm. (**B**) Representative Co-IF for PCDH20 (green), phalloidin (red) and DAPI (blue) in tuft cells from both wild-type and *Pcdh20-/-* small intestines. Scale bars, 5 μm. (**C**) SEM of a gallbladder tuft cell from a *Pcdh20+/+* mouse. Scale bar, 500 nm. (**D-E**) SEM of a gallbladder tuft cell from a *Pcdh20-/-* mouse highlighting the microvilli. Scale bar, 500 nm. Quantification of SEM images from *Pcdh20-/-* and control gallbladder tuft cells for (**F**) number of microvilli/area, (**G**) microvilli angle, (**H**) average vesicle size, and (**I**) number of vesicles. (**J**) Representative image illustrating the analysis pipeline for microvillar spacing using line transects and plot profile measurements. (**K-M**) Graphs showing spacing between microvilli at the top (**K**) middle (**L**) and bottom (**M**) of the microvillar apparatus. Data are represented as mean +/-SD. For immunostaining and SEM analyses, images from n = 3 *Pcdh20* +/+ and *Pcdh20* -/- mice were analyzed. ^*^, p < 0.05; ^**^, p < 0.001.

By SEM analysis, gallbladder tuft cells in wild-type (*Pcdh20+/+*) mice were found to be characterized by the tall, rigid microvilli previously described for this cell type (**Figure 6C**). In contrast, we found that tuft cell microvilli in the *Pcdh20*-/- model were stunted, disorganized, and splayed, losing their characteristic rigid, bundled appearance (**Figure 6D-E, Figure S9D-E**). *Pcdh20-/-* tuft cells had a similar number of microvilli per area as compared to *Pcdh20+/+* tuft cells (**Figure 6F**). To assess how well the microvilli were packed, we measured the angle of the microvilli; the closer to 180º, the tighter the structure and, hence, the more rigid. *Pcdh20+/+* microvilli were found to be straighter, or more vertical, and closer to a 180° angle than those in *Pcdh20-/-* tuft cells (**Figure 6G**). This value was significantly more variable in *Pcdh20-/-* tuft cells (3.18 vs. 1.01, F-test, p < 0.05). While modest, this deviation reflects a statistically significant loss of the highly organized, parallel architecture characteristic of the wild-type tuft, consistent with disrupted intermicrovillar linkages. Analysis of apical secretory vesicle size, for those budding from the tuft microvilli, trended toward being larger in *Pcdh20-/-* tuft cells, although this difference did not reach statistical significance (p = 0.1; **Figure 6H**). The number of vesicles, however, was significantly lower in the *Pcdh20-/-* tufts as compared to *Pcdh20+/+* tufts (**Figure 6I**). To assess microvillar spacing and organization, we drew linear transects at the top, middle, and bottom of each tuft cell and quantified peak-to-peak distances using the *Plot Profile* tool in Fiji (**Figure 6J**). Interestingly, spacing was significantly increased around the periphery of the *Pcdh20-/-* tuft cell microvillar tuft but remained comparable between genotypes in the central region (**Figure 6K-M**). Although the *mean* spacing did not differ significantly, the *Pcdh20-/-* tuft cells consistently displayed greater variance in microvillar spacing across all positions examined (top: 20,306 vs. 10,100.25; middle: 21,844.84 vs. 2,970.25; bottom: 15,376 vs. 3,821.71) reflecting disorder within the *Pcdh20-/-* tufts. Together with the observed defects in vesicle formation and the widely variable angles in the villi, these findings support a model in which PCDH20 serves as an important structural determinant of tuft cell microvillar organization.

### AlphaFold3 predicts that PCDH20 forms homodimer linkers

To structurally determine how PCDH20, CDHR2, and CDHR5 contribute to and reinforce tuft cell microvillar length and rigidity, we molecularly docked the relevant murine cadherins *in silico* using AlphaFold 3. We first tested whether AlphaFold could predict the interaction between CDHR2 and CDHR5 as this is a known interaction and part of the intermicrovillar attachment complex (IMAC)^22, 23^. When we docked CDHR2 and CDHR5 together, 4 of 5 computed models (80%) predicted an antiparallel interaction between the terminal two domains consistent with a cross-link between adjacent microvilli (**Figure 7A-B**). This interaction is consistent with a structural model created based on recombinantly expressed proteins.^59^

**Figure 7.**
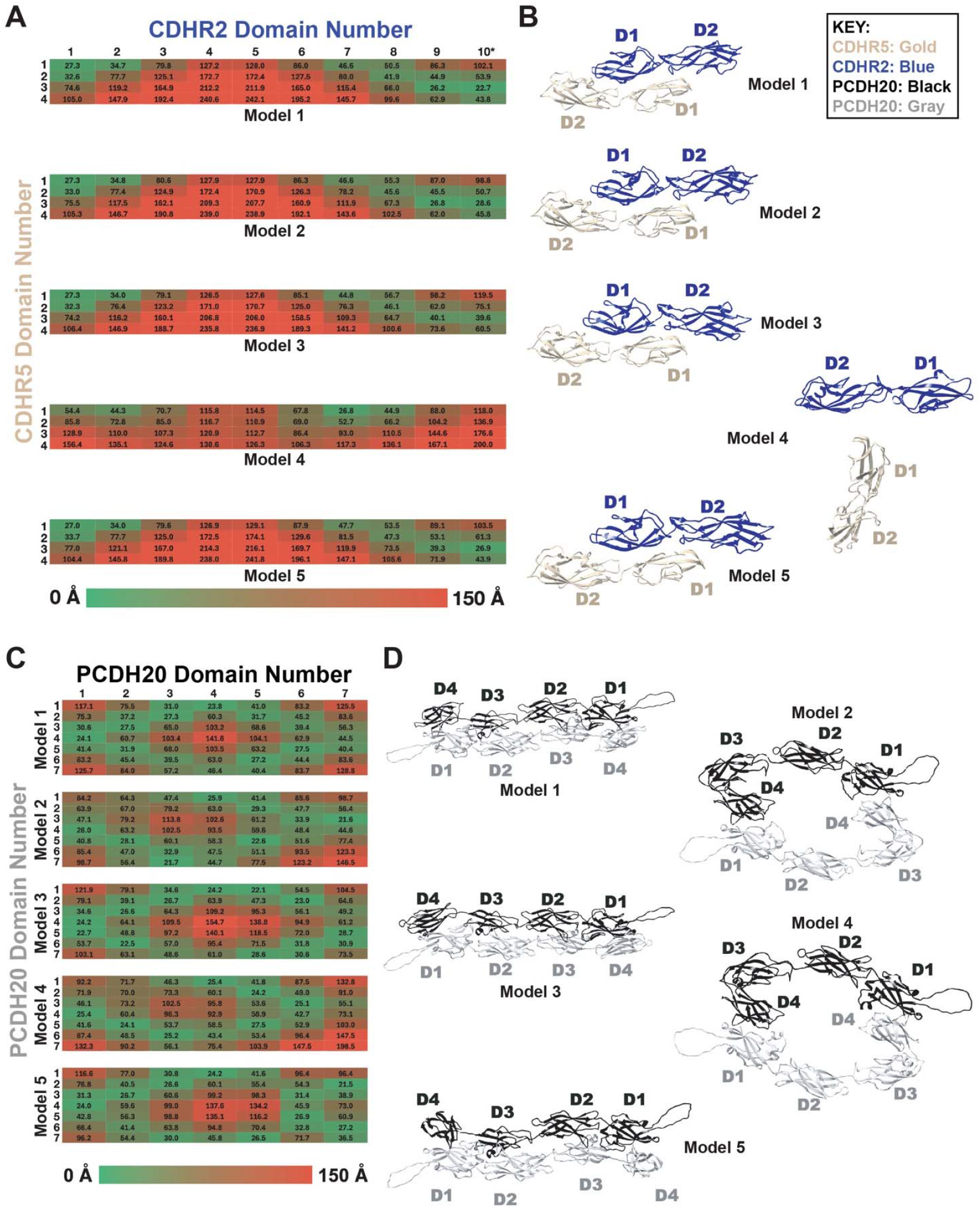
PCDH20 forms a homodimeric crosslink. (**A**) Heat map matrix displaying the center-to-center distance between each domain in CDHR2 (horizontal axis) and CDHR5 (vertical axis) for models 1 through 5. (**B**) Ribbon diagrams of the interactions between cadherin domains 1 and 2 of CDHR2 (blue) with cadherin domains 1 and 2 of CDHR5 (gold), displayed adjacent to the respective model. The remaining domains are omitted to not obscure important interactions. (**C**) Heat map matrix displaying the center-to-center distance between the domains of the first PCDH20 (horizontal axis) to the domains of the second PCDH20 (vertical axis) for models 1 through 5. (**D**) Ribbon diagrams of the interactions between cadherin domains 1-4 of the first PCDH20 (black) with those of the second PCDH20 protein (gray), displayed adjacent to the respective model. Remaining domains are omitted to not obscure important interactions. Heat maps are colored from green (0 Å) to red (150 Å).

With confidence in the ability of AlphaFold to dock cadherin domains, we next explored whether PCDH20 interacted with either CDHR5 or CDHR2 or both. All computed models between CDHR5 and PCDH20 yielded non-sensical interactions with domains being orthogonal to one another (**Figure S10A-B**). When CDHR2 was docked against PCDH20, however, AlphaFold3 suggested a parallel interaction (**Figure S10C-D)**. Such an interaction could allow lateral bundling of cadherins, a well described phenomenon; however, this would not serve as a structural cross-link between microvilli.

As heterodimeric docking between PCDH20 and either CDHR2 or CDHR5 did not yield plausible crosslinks, we next tested whether PCDH20 formed a homodimeric pair. All 5 of the predicted models of a PCDH20-PCDH20 homodimer (100%), suggested an interaction between the terminal domain 1 of PCDH20 with domain 4 of the second chain of PCDH20 (**Figure 7C-D**). Three of these models also predicted reciprocal interactions between domain 2 and domain 3 of the two PCDH20 chains. No intramolecular interactions were predicted. Such an antiparallel interaction between Domains 1-4 on each respective chain would yield a quaternary complex that could span and crosslink between two adjacent microvilli.

When the CDHR2:CDHR5 heterodimer is linearized, while maintaining the predicted interaction between domains 1 and 2 of each protein, a crosslinker length of ∼57 nm from outer membrane-to-outer membrane is obtained. A very similar length (∼55 nm) is obtained when the PCDH20 homodimer is linearized while maintaining the predicted interactions between domains 1-4 (**Figure 8A**). This predicted length is highly consistent with the experimentally measured distribution of intermicrovillar linkage lengths from our freeze etch EM data, which shows a peak around 50-60 nm (**Figure 8B**) Lastly, we modeled the CDHR2:CDHR5 heterodimer and the PCDH20 homodimer (**Figure 8C**) across adjacent microvilli to provide a visual representation of how these cadherin domains stabilize intermicrovillar interactions and why loss of PCDH20 results in loss of orientation and structural integrity even if the actin core bundle and myosin are intact.

**Figure 8.**
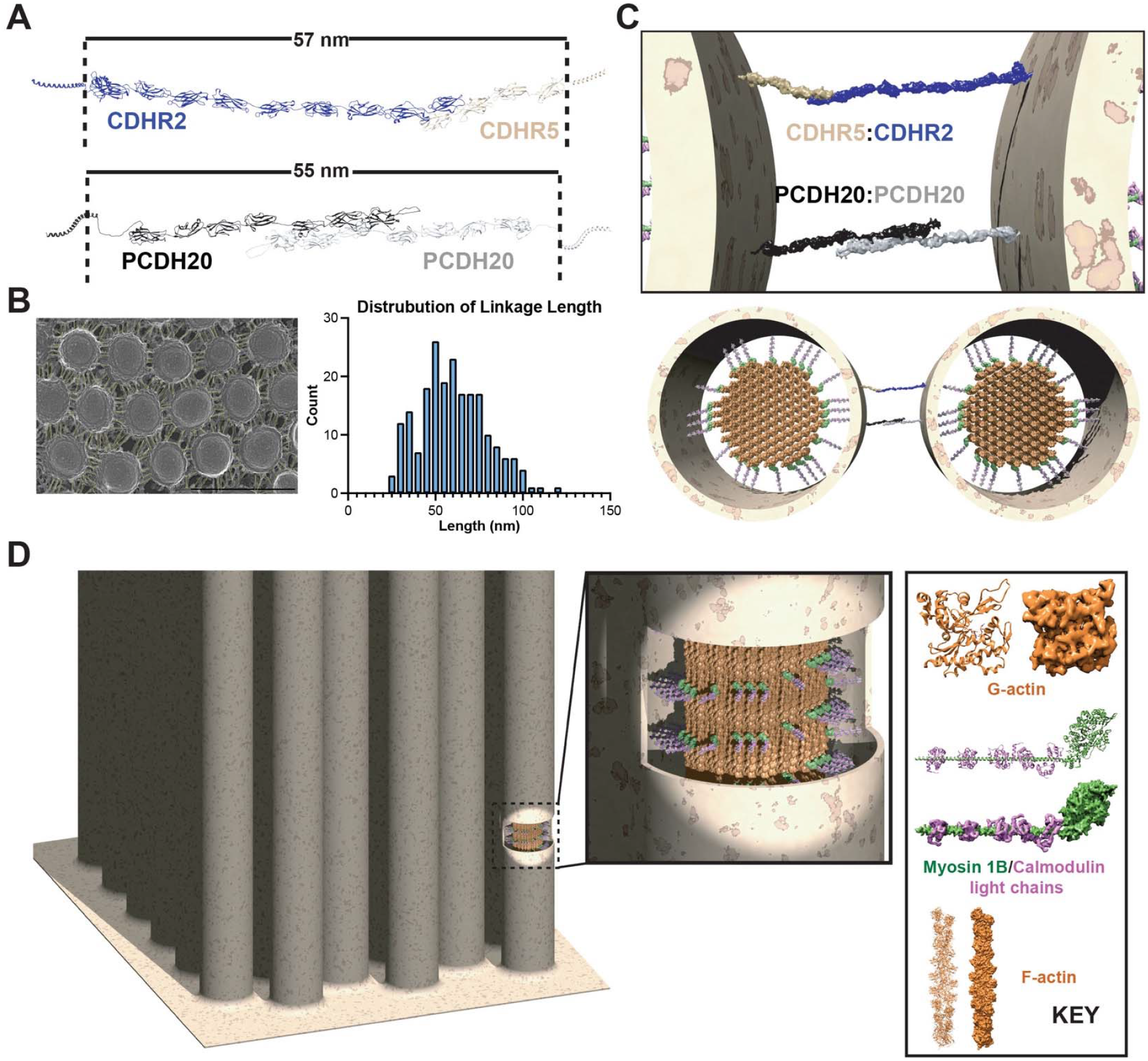
Molecular modeling of the tuft cell brush border. (**A**) *In silico* linearization of the CDHR2:CDHR5 heterodimer suggests a span of ∼57 nm, while linearization of the PCHD20 homodimer suggests a similar distance of ∼55 nm. This is consistent with (**B**) the average distance between tuft cell microvilli (∼ 50-60 nm) as measured from freeze etch electron microscopy. Scale bar, 500 nm. (**C**) Model of CDHR2:CDHR5 and PCDH20:PCDH20 demonstrating structural redundancy in intermicrovillar linkage. (**D**) Modeling of the actin core bundle decorated with Myosin 1B and calmodulin light chains spans the requisite ∼180 nm diameter to support the tuft cell microvillar membrane.

### Molecular Model of the Tuft Cell Microvillus

To help visualize the cadherin cross-links *in situ*, we created a molecular model of the tuft cell brush border. Each microvillus is supported by a core bundle of ∼100 actin filaments arranged in a hexagonal lattice (**Figure 8D, Figure S11**).^54^ The center-to-center spacing is 92 nm, which is much tighter than the 120 nm spacing of the actin core bundle in the enterocyte microvillus (**Figure S11**)^60-62^ Even with this narrow spacing, there is no steric hindrance between actin filaments as demonstrated in a clipping series orthogonal to the actin bundle (**Supplemental Movie 1**). However, such a narrow spacing will not accommodate regular crosslinks between actin bundling proteins like advillin and fimbrin/plastin (compare to the enterocyte microvillus model^60^) but would accommodate Espin (**Figure 4**)^63^. These proteins likely reside at regions of missing actin filament neighbors. Recent analysis of the tuft cell microvillar actin core bundle suggests that despite being hexagonally oriented, on average actin filaments only have 5 neighbors as opposed to the “ideal” 6 neighbors that we modeled in our saturated hexagonal lattice (**Figure S11**). The actin core bundle is laterally attached to the microvillar membrane by Myosin 1B (Myo1b). The structure of rat Myo1B has been reported, which allowed us to add this structure around the actin core bundle after extending the lever arm^64^. The size of the core bundle and Myo1b lateral crosslinks spans the ∼180 nm diameter of the tuft cell microvillus (**Figure 8C**). Altogether, the data presented here coupled with the existing literature allowed us to create a self-consistent molecular model of the cytoskeleton supporting individual tuft cell microvilli as well as the cadherin network responsible for organization of the apical microvillar network – the sine qua none attribute for which the tuft cell is named.

## DISCUSSION

Tuft cells are identified by their striking morphology; however, how form follows function for this cell type has yet to be determined. Though still a relatively young field, tuft cell biologists have identified roles for tuft cells in several disease states including infection (parasite, bacterial, viral), tissue injury, inflammation, and tumorigenesis^13^. A role for chemosensation has been demonstrated in several of these states, however whether tuft cells also signal via mechanotransduction remains a key unanswered question. Microvilli enhance membrane surface area, playing a role in absorption in enterocytes, however specialized microvilli, such as stereocilia, are required for critical mechanotransduction processes, such as in hearing and balance^20^. Because epithelial cell microvilli and inner ear hair cell stereocilia share many components, they are often compared to one another. Here, we showed that tuft cells share several important structural elements with cochlear hair cells. We identified common gene expression signatures including actin bundling, stereocilium assembly, and sensory perception of mechanical stimulus (**Figure 2**). These findings strongly support the long-standing hypothesis that tuft cells may possess mechanosensory capabilities. Our data suggest that targeted mutations known to cause deafness could also lead to tuft cell dysfunction, potentially manifesting as digestive or respiratory issues.

Among genes common to both our POU2F3 ChIP-seq analysis in tuft cells and the inner hair cell transcriptome, was *Pcdh20*. In hair cells, mechanoelectrical transduction is dependent on tip link proteins between stereocilia composed of PCDH15 and CDH23^20^. Based on our immunostaining and immunogold labeling studies localizing PCDH20 to tuft cell microvilli, we hypothesize that PCDH20 plays a similar role in tuft cells. Molecular modeling suggests that PCDH20 forms a homodimer as part of these intermicrovillar linkages (**Figure 7**), with potential to bind CDHR2 in a parallel fashion. The latter could result in a stronger link and more rigid microvilli (**Figure S10**). These models, however, require validation with biochemical assays and crystallography/electron microscopic reconstruction. Additional genes identified include several actin binding proteins, such as Espin, and other proteins involved in stereocilia elongation such as Whirlin, which are likely required to build the thick actin bundles characteristic of both cell types. Indeed, other recent studies showed both EPS8 and EPS8L2 localize to tuft cell microvilli tips^54^, and EPS8 interacts with whirlin at the tips of hair cell stereocilia^65, 66^. TMC1 – a major component of the hair cell stereocilia mechanotransduction channel - was identified in both the tuft cell and lung cancer cell POU2F3 ChIP-seq analyses, as well as in tuft cells at the protein level (**Figure 4**). TMC1 expression could indicate the presence of a tuft cell MET-like complex. Other possible MET components expressed at the mRNA level in tuft cells, but not regulated by POU2F3, include TMC4/5 and CIB2, which were recently described in intestinal enterocytes^67^. It is likely, though, that tuft cells will form complexes constructed of analogous, yet unique protein components as compared to other cell types.

Consistent with a role in intermicrovillar linkage, loss of PCDH20 resulted in abnormal tuft cell microvilli with a markedly splayed appearance in comparison to the normally tight bundle of microvilli in normal cells. We found gallbladder tuft cells in *Pcdh20-/-* mice to have irregular microvilli and fewer apical vesicles. These data suggest that loss of PCDH20 may interfere with apical tuft cell secretion. Loss of PCDH20 also significantly impacted the rigidity and perceived stiffness of the microvillar bundles. Molecular modeling suggests that these bundles are stabilized by both CDHR2:CDHR5 heterodimers and PCDH20 homodimers (**Figure 7-8**). This functional redundancy may explain why loss of PCDH20 did not result in a complete disarray of the microvillar tuft cell brush border.

Although a role for tuft cell microvilli in mechanotransduction has yet to be shown, loss of PCDH20 could potentially impact mechanotransduction in several ways: 1. By removing intermicrovillar linkages required to open TMC channels and 2. By influencing the amount of force sensed by the now stunted and/or collapsed microvilli. In the gallbladder, which changes dramatically in size based on an organism’s fed/fasted state, tuft cells may sense filling of the structure (during fasting), placing an applied force directly on the tuft. This could result in the release of known biliary tuft cell effectors, such as acetylcholine and leukotrienes^10^, to induce protective mucus release, thereby safeguarding against increasingly concentrated bile acids which are cytotoxic, and tonic gallbladder contraction which prevents bile stasis. In the intestines, tuft cells are subjected to luminal forces, which may include both applied force (filling, parasite infection) and a parallel, tensional force which would displace microvilli, resulting in release of multiple effectors (e.g. acetyl choline, IL-25). We hypothesize that loss of PCDH20 reduces the ability of tuft cells to sense these forces. Future experiments will assess whether or not tuft cell microvilli are mechanosensory; *in vitro* assays will be used to measure tuft cell microvillar stiffness and force sensing, while *in vivo* studies will define how PCDH20 loss alters tuft cell function in homeostatic and disease settings.

Our ChIP-seq analysis revealed that many structural proteins, including *Pcdh20*, are regulated by transcription factor POU2F3. This is consistent with the role of POU2F3 as a master regulator for tuft cell formation, dictating tuft cell fate, in part, by directing formation of the unique structure. Notably, ChIP-seq offers the advantage of detecting low-abundance transcripts which are often missed by conventional RNA-based approaches, allowing us to identify key tuft cell structural proteins even when their expression may be limited^68^. Many canonical tuft cell markers were identified in our analysis (e.g. *Il25, Ptgs1, Trpm5*, etc.) suggesting that POU2F3 directly drives expression of these genes. Interestingly, we also identified several neuronal genes in our ChIP-seq analysis. While many (e.g. DCLK1) have been described in tuft cells, it has also been shown that tuft and enteroendocrine cells are closely related in the intestines and diseased pancreas^69-72^. Loss of POU2F3 significantly increases abundance of enteroendocrine cells in pancreatic tumorigenesis suggesting that it not only drives expression of tuft cell genes but may also inhibit enteroendocrine-specific gene expression during tuft cell formation^73^. Future studies will focus on how POU2F3 both positively and negatively regulates tuft cell fate and identifying any critical binding partners (SOX9, RUNX1, etc.)

Altogether, this study identifies *Pcdh20* as a POU2F3 target gene and critical structural protein in tuft cells. To our knowledge, this is the first study demonstrating impaired tuft cell formation in response to structural protein loss. These data will serve as foundation for future studies on structure-function relationships in tuft cells.

## MATERIALS AND METHODS

All authors had access to the study data and reviewed and approved the final manuscript.

### Mice

Mice were housed in accordance with National Institutes of Health guidelines in American Association for Accreditation of Laboratory Animal Care-accredited facilities at Vanderbilt University, The Salk Institute for Biological Studies, the University of California San Diego, the University of Virginia, and the National Institutes of Health. The animal holding rooms were maintained at 70°F +/-2 degrees. Animal rooms are pathogen free, excluding common endemic pathogens. Wild type male C57B6/J mice were purchased from Jackson Laboratories. *Pcdh20-/-* mice were generated using clustered regularly interspaced short palindromic repeats (CRISPR). The gRNA to the mouse PCDH20 gene and Cas9 mRNA were co-injected into fertilized mouse eggs to generate targeted knockout offspring. F0 founder animals were identified by PCR product sequence analysis and were confirmed by off-target analysis. PCDH20 knockdown mice were genotyped using the following primers which recognize both wild-type (374 base pairs) and mutant alleles (323 base pairs): mPCDH20-5b, 5’-GGA AGT TTA GCA TTG TCC CTG G-3’, mPCDH20-3b, 5’-GGT TGT TTA GGG TCA CAT ACT GG-3’.

### Tuft cell preparation and isolation

Intestinal tuft cells were isolated as previously described^5, 12^. The proximal 5 cm of the murine small intestine was first dissected, then flushed, cut longitudinally, and rinsed to remove luminal contents. Tissues were incubated in a 37°C shaker for 20 minutes, in 5 mL 1 x PBS containing 2.5 mmol/L EDTA, 0.75 mmol/L dithiothreitol, and 10 mg/mL DNAse I. Tissues were shaken vigorously for 30 seconds, large pieces of intestinal wall were removed, and cells were spun down at 4°C, 1200 revolutions/minute, for 5 minutes. Supernatant was removed, and cells were resuspended in 5 mL Hank’s balanced salt solution with 1.0 U/mL dispase and 10mg/mL DNAse I, shaking for 10 minutes at 37°C. Digested cells were passed through a 100 μm filter and incubated with ACK lysing buffer. Single-cell suspensions were incubated on ice with mouse Fc receptor block (BD Biosciences, San Jose, CA; 1:200) followed by antigen-specific antibodies in FACS buffer. Then, 40,6-diamidino-2-phenylindole (Molecular Probes, Eugene, OR; 1:1000) was used to exclude dead cells. Cells were labeled with CD45 (Alexa Fluor 488), EpCAM (Alexa Fluor 647), and Siglec F (PE) (all BioLegend, 1:200). Fluorescence-minus-1 staining controls were included for gating populations of interest. Cells were FACS purified at the Salk Institute’s Flow Cytometry core facility on a BD Biosciences Influx cell sorter (100 μm size nozzle, 1 x PBS sheath buffer, with sheath pressure set to 20 pounds/square inch) or an Aria Fusion cell sorter (BD Biosciences Billerica, MA). Cells were sorted in 1-drop single-cell sort mode for counting accuracy into DMEM + 10% FBS.

### POU2F3 chromatin immunoprecipitation (ChIP) sequencing

Due to the low abundance of tuft cells in the murine small intestines (∼0.4%)^5^, we adapted a low input ChIP protocol called ChIPmentation with minor modifications^74^. Two pools of tuft cells from four adult male C57BL6/J mice were used as biological replicates. Approximately 1x10^4^ sorted tuft cells were crosslinked with 1% PFA at room temperature for 10 minutes, sonicated with Covaris M220 into 200-700 bp fragments, and incubated with 0.6 μg of anti-POU2F3 antibody (Santa Cruz #SC330) or control antibody (anti-IgG Cell Signaling #2729) overnight at 4°C. Protein A bead pull down (Invitrogen 10001D), washing, on-beads tagmentation (Illumina Nextera FC-121-1030), reverse crosslinking, library amplification (16-18 PCR cycles) and DNA purification were performed as described. The input library was prepared by tagmentation of 1 ng of sonicated and reverse crosslinked mouse chromatin and then amplified and processed as the ChIP DNA. The control and ChIP samples were analyzed by Agilent TapeStation, and 50 bp single-end sequencing was performed with Illumina HiSeq 2500.

### POU2F3 ChIP-seq analysis and comparison to hair cells

ChIP-seq data analyses were performed as previously described^75^. In brief, sequencing reads were checked by FastQC and aligned to mm9 with Bowtie (-k 1 -m 20 --best -S -n 2 -l 40)^76^. MACS2 (--nomodel --shift 100 --bw 150 --SPMR) was used to remove duplicated reads, call peaks and generate Bedgraph files that show fold change enrichment over input^77^. To ensure the quality of the POU2F3 binding sites, only peaks common between the two biological replicates (FDR<0.05) were called, and any peaks that overlap with the control IgG peaks (FDR<0.05) and the Encode blacklist regions were removed. Bedgraph files were then converted into BigWig files and uploaded to the UCSC Genome Browser for visualization. Signal profiling and correlation analysis were performed using deepTools^78^. Functional annotation of peaks and peak-gene association were done with GREAT (version 3.0.0) using the default “basal plus extension” parameter^79^. HOMER was used for motif enrichment analysis with size setting to 1000 bps^80^. Signals are visualized with trackplot R package (https://github.com/PoisonAlien/trackplot) for POU2F3 ChIP and IgG control across different gene regions.

### Comparison to tuft cell and hair cell RNA-seq datasets

The differentially expressed genes in mouse inner hair cells (P20) from a publicly available dataset^37^ were compared with POU2F3 ChIP-seq target genes. GO Enrichment Analysis was performed on the overlapping gene set using the enrichGO function from the clusterProfiler R package (https://github.com/YuLab-SMU/clusterProfiler).

### Computational Analysis of RNA-seq datasets

Single-cell RNA sequencing (scRNA-seq) data generated from gallbladder and extrahepatic duct tissue from *Pou2f3+/+* and *Pou2f3*-/- mice^11^ (GEO accession: GSE194035) were obtained from the GEO database in filtered_feature_bc_matrix.h5 format. Data processing was performed using the Seurat V5.1^81^ in R. For quality control, only cells with a minimum of 200 detected genes and present in at least 3 cells were preserved. Cells were then filtered to retain those with 300–5000 features, <20,000 RNA counts, <15% mitochondrial genes, and <0.1% hemoglobin expression. Following QC, the datasets were merged. Data normalization and variance stabilization were then performed using SCTransform while regressing out mitochondrial gene expression. Dimensionality reduction was achieved via principal component analysis (PCA), and cell-to-cell distances were computed using FindNeighbors (dims 1:50). Unsupervised clustering was performed with a resolution of 0.6 (with 30 starts), followed by Uniform Manifold Approximation and Projection (UMAP) for visualization of the cellular landscape. Feature plots for selected genes of interest (including *Pcdh20, Whrn, Espn*, and *Clic5*) were generated using Featureplot() based on orig.ident and arranged in a combined layout for comparison between the two genotypes.

### Fluorescence Microscopy and quantification

Fluorescence microscopy was performed as previously described^12, 31^. Briefly, tissues were fixed overnight in 4% paraformaldehyde, washed 3 times with PBS, and floated overnight in 30% sucrose at 4°C. Tissues were then incubated in a 1:1 mixture of 30% sucrose and Tissue-Tek optimal cutting temperature compound (OCT) (VWR, Radnor, PA) for 30 minutes, embedded in OCT, and frozen at -80°C. 7-10 μm tissue sections were cut, permeabilized with 0.1% Triton X-100 in 1x PBS, and blocked with 5% normal donkey serum and 1% BSA in PBS for 1 hour at room temperature. Tissue sections were stained with primary antibodies (Table S1) in 1 x PBS supplemented with 1% BSA and 0.1% Triton X-100 overnight. In some experiments, a polyclonal antibody against PCDH20, made against C-terminal amino acid 940-952, CMRERKPVDISNI, of the mouse protein (AB_3676540) was used^82^. Sections were then washed for 3 x 15 minutes in PBS with 1% Triton X-100, incubated with Alexa Fluor secondary antibodies and/or phalloidin (Invitrogen, Waltham, MA), washed again for 3 x 5 minutes, rinsed with distilled water, and mounted with Prolong Gold containing 4’,6-diamidino-2-phenylindole (Thermo Fisher). Tissues were imaged on either a Zeiss (Oberkochen, Germany) 710 confocal microscope, a Zeiss 880 or 980 Airyscan Super-Resolution microscope, an Andor BC43, or a Leica (Wetzlar, Germany) MICA confocal microscope using an HC PL APO 20x/0.75 or HC PL APO 63x/1.20 W objective. For quantification in FIJI, tuft cells were identified by DCLK1 staining and phalloidin boundaries, tuft cell ROIs were drawn based on those boundaries, and both total tuft cell area as well as area of PCDH20+ signal was measured within the tuft cell ROI in FIJI. Data was plotted as % of PCDH20+ area per tuft cell area compiled into Prism Graphpad for visualization and statistical analysis.

### Whole Mount Immunofluorescence

Tissues were dissected from wild-type mice immediately after euthanasia and cut into 0.5 cm X 0.5 cm squares. Tissues were fixed in 4 % PFA in PBS for 1 hour before washing in PBS 3 x 15 minutes. Tissues were then incubated 1:1000 Alexa Fluor 405 Phalloidin in 0.5 % PBST for 30 minutes, before washing in PBS for 15 minutes. Tissues were blocked by incubating in 10% NGS for 4 hours, before incubating in 1:400 primary antibody overnight at 4 C. Tissues were then washed again in PBS 3 x 15 minutes before incubating in 1:1000 anti-rabbit secondary antibody for 1 hour. Finally, tissues were washed again in PBS 3 x 15 minutes before mounting on slides with ProLong Gold antifade mounting media. TMC1 images were captured on a Nikon ECLIPSE Ti2-E inverted microscope with Yokogawa CSU-W1 Spinning Disk and Hamamatsu ORCA-Fusion BT Digital CMOS camera C15440-20UP.

For inner ear immunolabeling, 1 mL of the above-described fixative was injected using an insulin syringe into the round window of freshly dissected WT and PCDH20 KO mice inner ears and left for 30 min into a glass vial with another 10 mL of fixative at RT. Then inner ears were PBS washed (3 x 15 min), carefully dissected under a dissecting scope, and transferred to a glass plate (Pyrex® spot plate with nine depressions). Tissues were permeabilized with 0.05% triton-X (10 min), blocked with 10% NGS (1h), incubated with chicken anti-PCDH20 primary antibody 1:200 (overnight 4^°^C), washed with PBS + 1% NGS (3 x 10 min), incubated with anti-chicken secondary antibody 1:500 and phalloidin Alexa-546 1:500 (40 min RT, Invitrogen), washed in PBS, mounted with ProLong Glass antifade (Invitrogen) and imaged in a Zeiss 880 Airyscan microscope with a 63x 1.4NA oil objective with a 42nm xy-pixel size and a 110nm z-step spacing, then 3D Airyscan processed with automatic settings.

### PCDH20 Transfection

HeLa or U2OS cells were passaged in DMEM supplemented with 10% FBS and 100 μg/mL penicillin-streptomycin and were split at a ratio of 1:10 once every two to three days using Trypsin. At 80% confluency, cells were transfected with either GFP or PCDH20-GFP (generated by the Gong Laboratory). 3.7 μg of PCDH20 plasmid was diluted 1:1000 in OptiMeM and cells were transfected using Lipofectamine for 48 hrs. Cells were then washed and fixed with paraformaldehyde, stained, and imaged as previously described.

### Scanning Electron Microscopy (SEM)

SEM was performed as previously described^13, 31^. Briefly, murine small intestines or gallbladders from wild-type or PCDH20 knockdown C57Bl6/J mice were freshly dissected, cut into 2-mm fragments and immediately fixed with 4% formaldehyde, 2.5% glutaraldehyde, 0.1M sodium cacodylate buffer (pH 7.2), and 2mM CaCl2 for 2 hours at room temperature. Samples were then washed for 3 x 15 minutes in the same buffer and incubated with 2 consecutive baths of 1% osmium tetroxide and 1% tannic acid for 1 hour each with 3 x□10-minute water washes in between each bath. Samples were dehydrated in a graded series of ethanols until absolute and critical point dried (Leica CPD300 or Tousimi PVT 3D). All reagents were bought from Electron Microscopy Sciences, Hatfield, PA. They were then mounted on carbon tape covering aluminum stubs and sputtered coated with platinum (4 nm) (Leica EM SCD500 or Cressington 108). Tissues were imaged by FEG-SEM using a Zeiss Sigma VP operated at 5 kV or a Zeiss Crossbeam 550 operated at 2 kV and were photographed at 2048□x 1536 pixels of resolution. *SEM Analysis*. SEM images were analyzed using FIJI^83^ and Graphpad Prism. Briefly, the scale was set for each individual image. Microvillar angles were measured on tuft projections with FIJI’s angle tool. Vesicles were drawn manually on all microvillar projections in each tuft; ROIs were filled in and quantified using the Analyze Particles function. To analyze microvillar spacing, lines were drawn at the top, middle, and bottom of the microvillar apparatus, and the *Plot Profiles* tool was used. The spacing between microvilli was then measured. Data was compiled into Prism Graphpad for visualization and statistical analysis.

### Scanning EM of ultrathin sections

Murine small intestines were processed and imaged as described previously with some modifications^84^. Materials were sourced from Electron Microscopy Sciences (Hatfield, PA) unless noted otherwise. Briefly, fresh biopsies of small intestines were dissected into small pieces (<1 mm) and quickly immersed in 37°C buffered fixative (2.5% glutaraldehyde, 2% paraformaldehyde, 3 mM CaCl2, 0.1M sodium cacodylate) before overnight storage at 4°C in the same solution^85^. Samples were serially rinsed with ice-cold cacodylate/CaCl2 buffer and stained with buffered reduced osmium (1.5% Osmium, 1.5% potassium ferrocyanide, 3 mM CaCl2, 0.1M sodium cacodylate) for 45 minutes in the dark at room temperature. Samples were serially rinsed with ice-cold distilled water before 30 minutes of treatment with filtered 1% aqueous thiocarbohydrazide (Ted Pella, Redding, CA) that was prepared at 60°C for an hour before use. Samples were serially rinsed with ice-cold distilled water before staining with 1.5% aqueous osmium tetroxide for 45 minutes in the dark at room temperature. Samples were rinsed serially with ice-cold water before staining overnight with 1% uranyl acetate at 4°C. Samples were then rinsed with distilled water and stained with Walton’s lead aspartate for 30 minutes at 60°C^86^. Samples were rinsed again with distilled water and serially dehydrated in ice cold ethanol. Samples were then infiltrated with Durcupan resin (hard formulation) and polymerized at 60°C for 3 days. A sample block was prepared for serial sectioning as previously described and a ribbon of 248 serial sections, each of dimension ∼150 μm x 400 μm x 70 nm (x-y-z), were collected onto a silicon wafer diced to a dimension of 35 x 6 mm using Diatome diamond knives mounted in a Leica UC7 ultramicrotome^87^. The chip was imaged using a Zeiss Sigma VP scanning electron microscope equipped with a Gatan backscattered electron detector and Atlas5 (Fibics) control software. Images were collected at a working distance of between 4-5mm and with an accelerating voltage of 3kV and pixel sizes between 2-8nm.

### Freeze fracture and etching Electron Microscopy (FFE-EM)

Preparation of fixed samples: Samples were incubated with 2% glutaraldehyde overnight and then washed in ddH_2_O. Fixed samples were then fast frozen using a Life Cell CF-100 slam-freezing machine. Frozen tissue was subjected to freeze-fracture at -110°C in a Balzers freeze-fracture machine, followed by freeze-etch at -100°C for 10 min. Freeze-etched samples were rotary shadowed with platinum and stabilized with carbon using electron-beam metal-evaporation guns (Cressington Scientific) to create replicas of the exposed surfaces. Replicas were cleaned with sodium hypochlorite, washed with ddH_2_O, and transferred onto 300 mesh hexagonal copper grids (Electron Microscopy Sciences). Replicas were imaged on a JEOL 2100 transmission electron microscope with an Orius 832 CCD camera (Gatan). Images were captured with DigitalMicrograph (Gatan).

### Transmission Electron Microscopy and Immunogold Labeling of Ultrathin Sections and Imaging

Gallbladder was fixed in 0.5% glutaraldehyde, 4% paraformaldehyde in 0.1 M sodium cacodylate for 1 hour then gradually cryoprotected to a final concentration of 30% glycerol in 50 mM HEPES. Samples were plunge frozen in liquid ethane. Frozen samples were freeze-substituted in a Leica Biosystems AFS2 with 1.5% uranyl acetate in absolute methanol at -90°C for 2 days, then infiltrated with HM20 Lowicryl resin (Electron Microscopy Sciences) over the course of 2 days at -45°C. Finally, the resin was UV-polymerized for 2 days between -45°C and 0°C. Ultrathin sections (70 to 100 nm) were cut and collected on 300-mesh Ni grids (Electron Microscopy Sciences). Grids were post-stained with 2% uranyl acetate and lead citrate. Samples were imaged on a 200-a JEOL 2100 Plus at 200 keV equipped with an AMT nanosprint15mkII CMOS camera; processing was done with FIJI and was limited to cropping and linear adjustments to brightness and contrast. Immunogold labeling was done like previously described^63^. In brief, samples were antigen retrieved using 0.1% sodium borohydride, 50 mM glycine in tris buffered salt solution (TBST, 5 mM Tris, 0.81% NaCl, 0.1% Triton X-100) for 10 minutes followed by blocking in 10% normal goat serum in TBST for 30 minutes. Samples were stained with primary antibody (PCDH20, Gong Laboratory) at a dilution of 1:100 in TBST with 1% normal goat serum for 1 hour. Samples were extensively washed in TBST with 1% normal goat serum and then stained with 10 nm colloidal gold conjugated to goat anti-chicken secondary (Electron Microscopy Sciences) for 1 hour. Samples were washed in TBST, followed by ddH_2_O. Grids were post stained with 2% uranyl acetate. Immunogold labeled samples were imaged on a JEOL 2100 Plus transmission electron microscope with an AMT nanosprint15mkII camera using SerialEM to acquire the images.

### Molecular Docking

Potential homo- and heterodimeric interactions between CDHR2, CDHR5, and PCDH20 were identified by uploading the murine peptide sequences into Alphafold3 web server^88^ (on June 5^th^, 2025), which generates five dimeric conformations. The proximity between cadherin domains were determined by measuring the distance (equation 1) between the center of mass of each cadherin domain to one another as calculated with ChimeraX molecular graphics program^89-92^. After identifying the cadherin domains predicted to associate with one another, the structures were inspected to determine whether the dimeric interactions were compatible with the known orientation of cadherins (e.g. parallel or antiparallel) or whether the interactions where non-sensical (e.g. orthogonal). The structures were then linearized using ChimeraX using the match command and sequences threaded with the program modeller 10.7^93, 94^.

Equation 1. Distance (d) calculated between the center of mass of two cadherin domains (x_1_,y_1_,z_1_) (x_2_,y_2_,z_2_) in 3-Dimensional Cartesian coordinates.

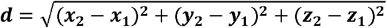

### Molecular Modeling

The molecular structure of the actin filament was created from the coordinates and rotational/translational parameters of actin as determined by X-ray fiber diffraction (PDB ID: 2ZWH)^95^. In an identical fashion and using the same scripts as we have previously reported when we generated a molecular model of the enterocyte brush border^60^. A molecular model for the F-actin bundle was created by modifying the hexagonal lattice scripts used for the enterocyte brush border to shorten the interfilament distances from 120 to 92 Angstroms and expand the hexagonal lattice from 19 to 109 filaments consistent with the data reported here and by Tyska’s group^55^. Myosin 1B was modeled using the cryoEM structures or rat Myosin-1B bound to actin (PDB IDs: 5V7X, 6C1D, 6C1G, and 6C1H)^64^. Due to structural flexibility in myosin lever-arm/light-chain binding domain coordinates were not present in the deposited structures. These were modeled by extending the truncated alpha helical lever arm in ChimeraX^92^ and adding calmodulin domains to the five IQ domains as we have done previously with myosin 1A^60^. Myosins were placed outwardly facing around the outer ring of the actin bundle in accord with the established actin-myosin 1B interaction structure^64^. The structure and molecular surfaces were created with UCSF chimera^91^ and exported in POV-Ray format (1 angstrom = 1 POV-Ray unit) for additional features to be added. The tuft cell membrane was added in POV-Ray and rendered as we have described previously^60, 96, 97^.

### Statistical Analysis

Statistical analyses and data processing were performed in Image J or Graphpad Prism (GraphPad, San Diego, CA). Statistical significance was calculated by either 2-tailed unpaired t-tests assuming equal variance, a t-test with a Welch’s correction (if variances were unequal), a Mann-Whitney test (if the data was non-normally distributed) or 1-way analysis of variance. Data are expressed as mean ± standard deviation. Figures were generated with Adobe Photoshop and Illustrator. Figure 7 was generated with bioRender.

## Abbreviations

CDH23: cadherin 23
ChIP-seq: chromatin immunoprecipitation sequencing
CRISPR: clustered regularly interspaced short palindromic repeats
EM: electron microscopy
FACS: fluorescence activated cell sorting
FFE-EM: freeze fracture deep etching electron microscopy
GEMM: genetically engineered mouse models
GO: Gene Ontology
GREAT: Genomic Regions Enrichment of Annotations
GWAS: Genome-wide Association Analysis
IF: immunofluorescence
MET: mechanotransducer non-selective cation channel complex
mRNA: messenger RNA
NSCLC: non-small cell lung cancer
PCA: principal component analysis
PCDH15: protocadherin 15
PCDH20: protocadherin 20
POU2F3: POU Class 2 Homeobox 3
SEM: scanning electron microscopy
TEM: transmission electron microscopy
TMC1: transmembrane channel-like protein 1
TMC2: transmembrane channel-like protein 2
TSS: transcription start site

## ACKNOWLEDGEMENTS

The authors would like to thank Ian Junker and Amelia Cephas for technical assistance, Jakob von Moltke for supplying tuft cell EM images, and Claire O’Leary for supplying gallbladder scRNA-seq data and for critical reading of the manuscript. This work was supported by the Waitt Advanced Biophotonics Core Facility of the Salk Institute (RRID:SCR_014838) with funding from NIH-NCI CCSG P30 CA014195, NIH-NIA San Diego Nathan Shock Center P30 AG068635, The Henry L. Guenther Foundation and the Waitt Foundation.

---

**Table S1.** Antibodies used in study. IF, immunofluorescence; ChIP-seq, chromatin immunoprecipitation sequencing.

| Antibody | Provided by | Catalog # | Species | Dilution, ChIP-seq, IF, flow cytometry |
| --- | --- | --- | --- | --- |
| CD45, Alexa Fluor 488 | Biolegend | 103122 | Rat | 1:200, Flow Cytometry |
| DCLK1 | Abcam | 31704 | Rabbit | 1:1000, IF |
| EpCAM, Alexa Fluor 647 | Biolegend | 118212 | Rat | 1:200, Flow Cytometry |
| Anti-IgG | Cell Signaling | 2729 | Rabbit | 0.6 µg, ChIP-seq |
| Pan-Espin | Dr. Kachar | N/A | Rabbit | 1:250, IF |
| PCDH20 | Novus | NBP2-30010 | Rabbit | 1:500 – 1:1000, IF |
| PCDH20 | Dr. Gong | N/A | Chicken | 1:1000, IF |
| Phalloidin | Invitrogen | A22287,<br>A12381,<br>A30104 | N/A | 1:500, IF |
| POU2F3 | Santa Cruz | SC-330 | Rabbit | 0.6 µg, ChIP-seq |
| Siglec F, PE | Biolegend | 155506 | Rat | 1:200, Flow Cytometry |
| Siglec F | BD Biosciences | 552125 | Rat | 1:200 IF |
| TMC1 | Sigma | ABN1649M | Rabbit | 1:400, IF |
| Whirlin | Dr. Kachar | N/A | Rabbit | 1:250, IF |

**Figure S1.**
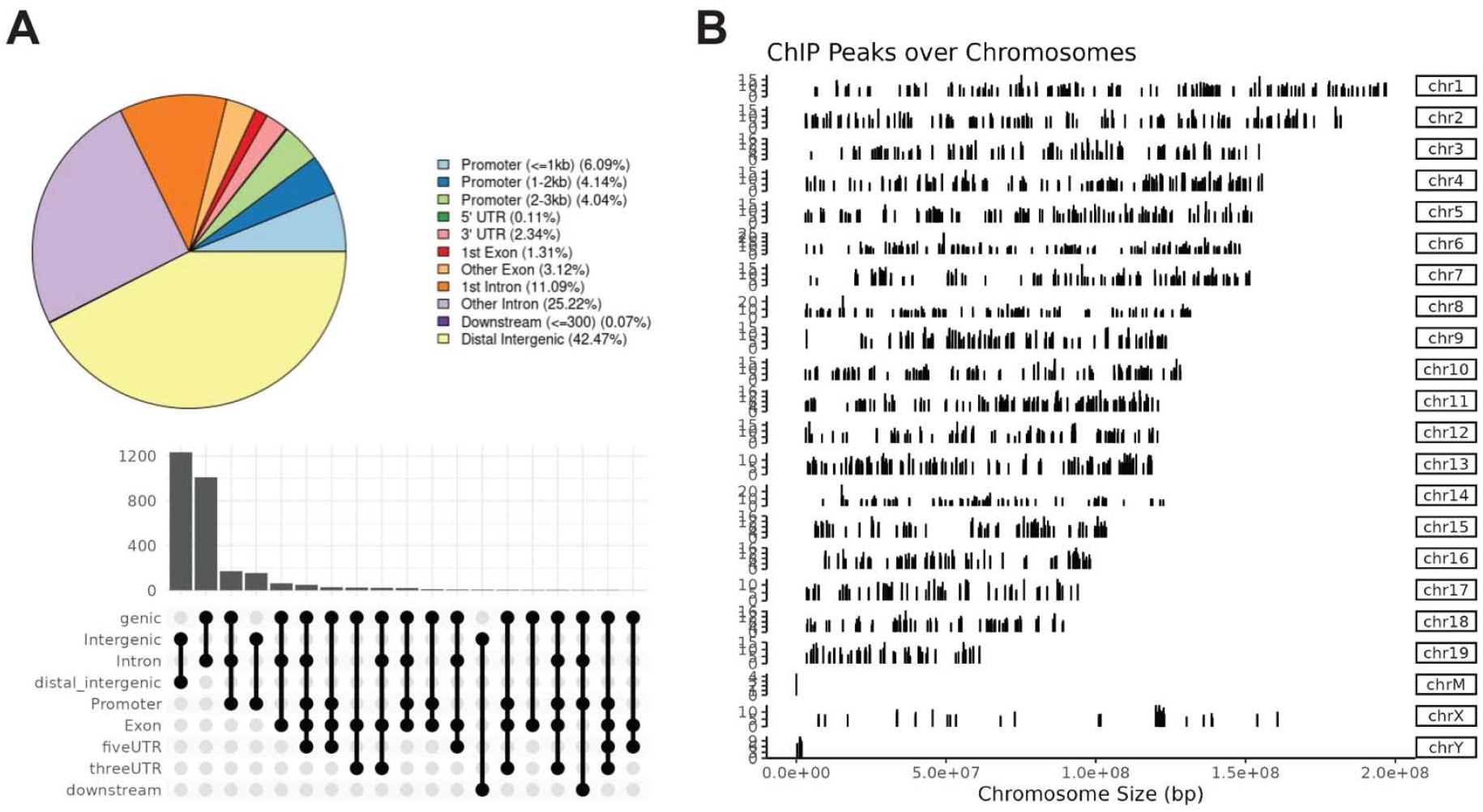
ChIP-seq identifies POU2F3 target genes in tuft cells. (**A**) Location of POU2F3 binding sites within target genes. (**B**) Distribution of POU2F3 binding sites throughout the murine genome by chromosome.

**Figure S2.**
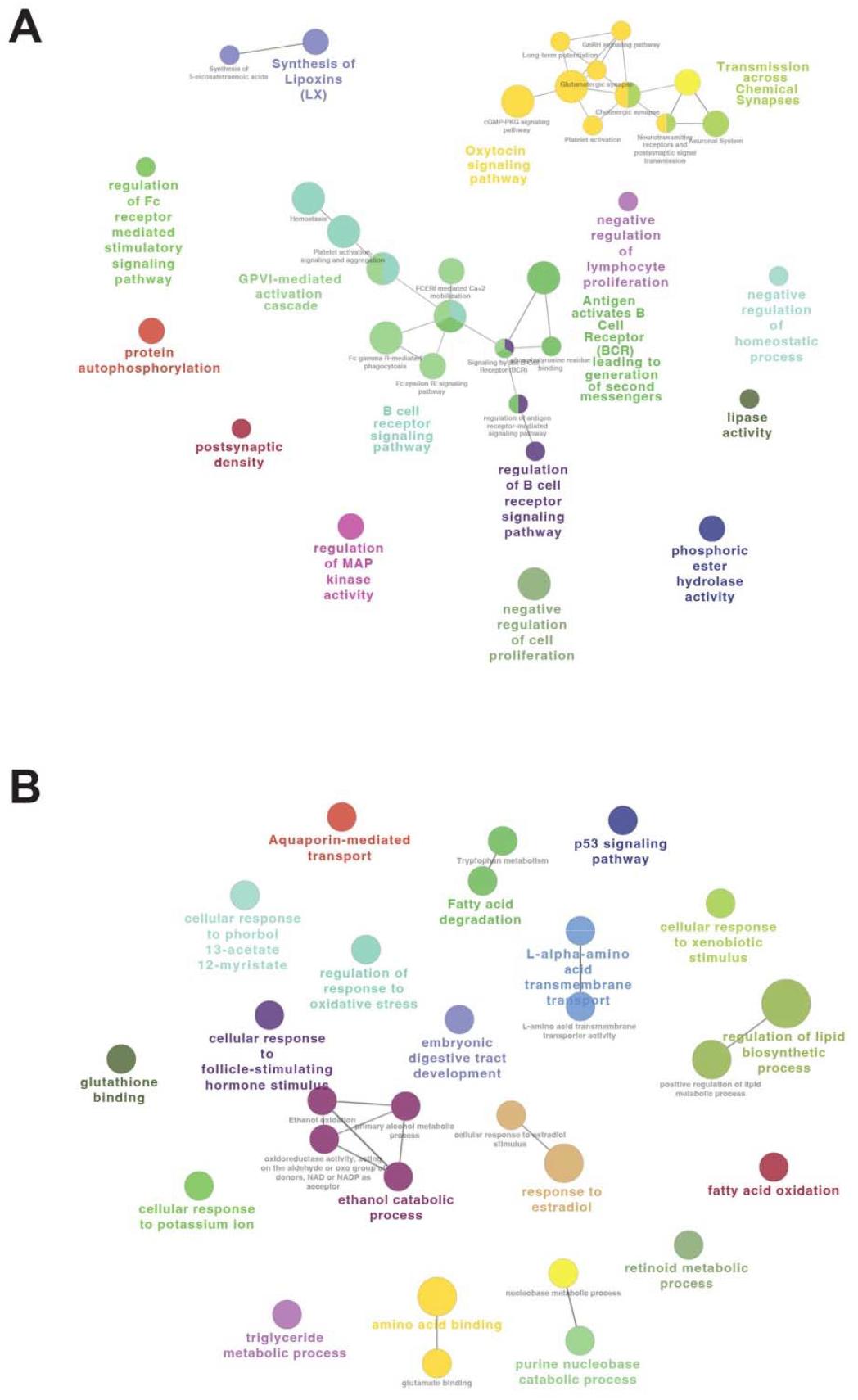
Gene Ontology (GO) identifies predicted pathways regulated by transcription factor POU2F3 in tuft cells. Biological role of genes common to the POU2F3-ChIP-seq dataset and (**A**) genes differentially upregulated in duodenal tuft cells (350 genes) or (**B**) differentially downregulated (120 genes), visualized with ClueGO. Nodes represent enriched GO terms and node size reflects statistical significance. Edges reflect the connectivity between GO terms (kappa statistics). GO terms are grouped into functional groups by color based on shared genes and Cohen kappa coefficient statistics.

**Figure S3.**
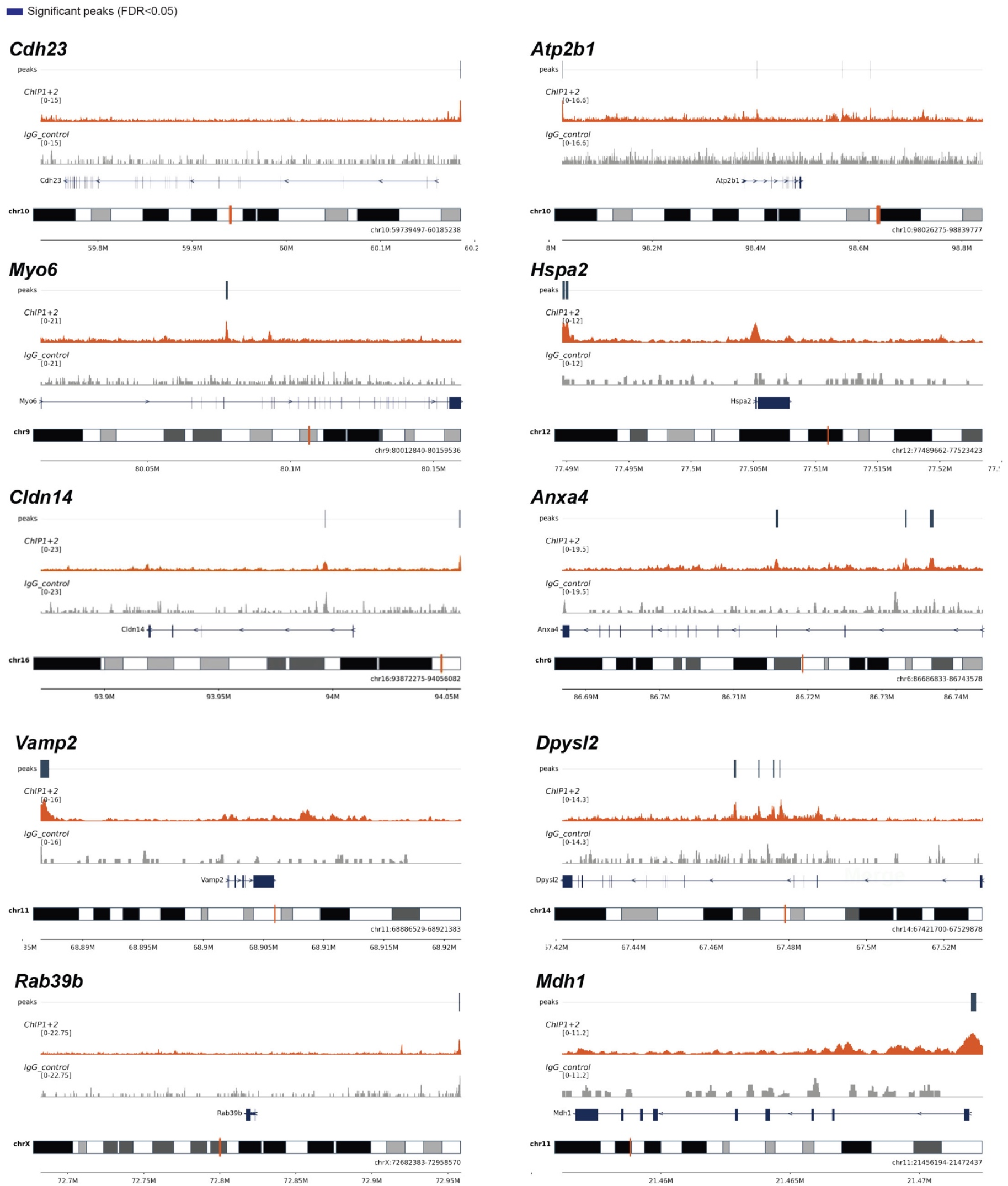
POU2F3 ChIP-seq reveals a cochlear hair cell gene signature. Tracks of select genes common to both the POU2F3 ChIP-seq analysis performed on duodenal tuft cells and a scRNA-seq dataset generated from adult murine cochlear inner hair cells (Figure 2) from a study by Jean et al.

**Figure S4.**
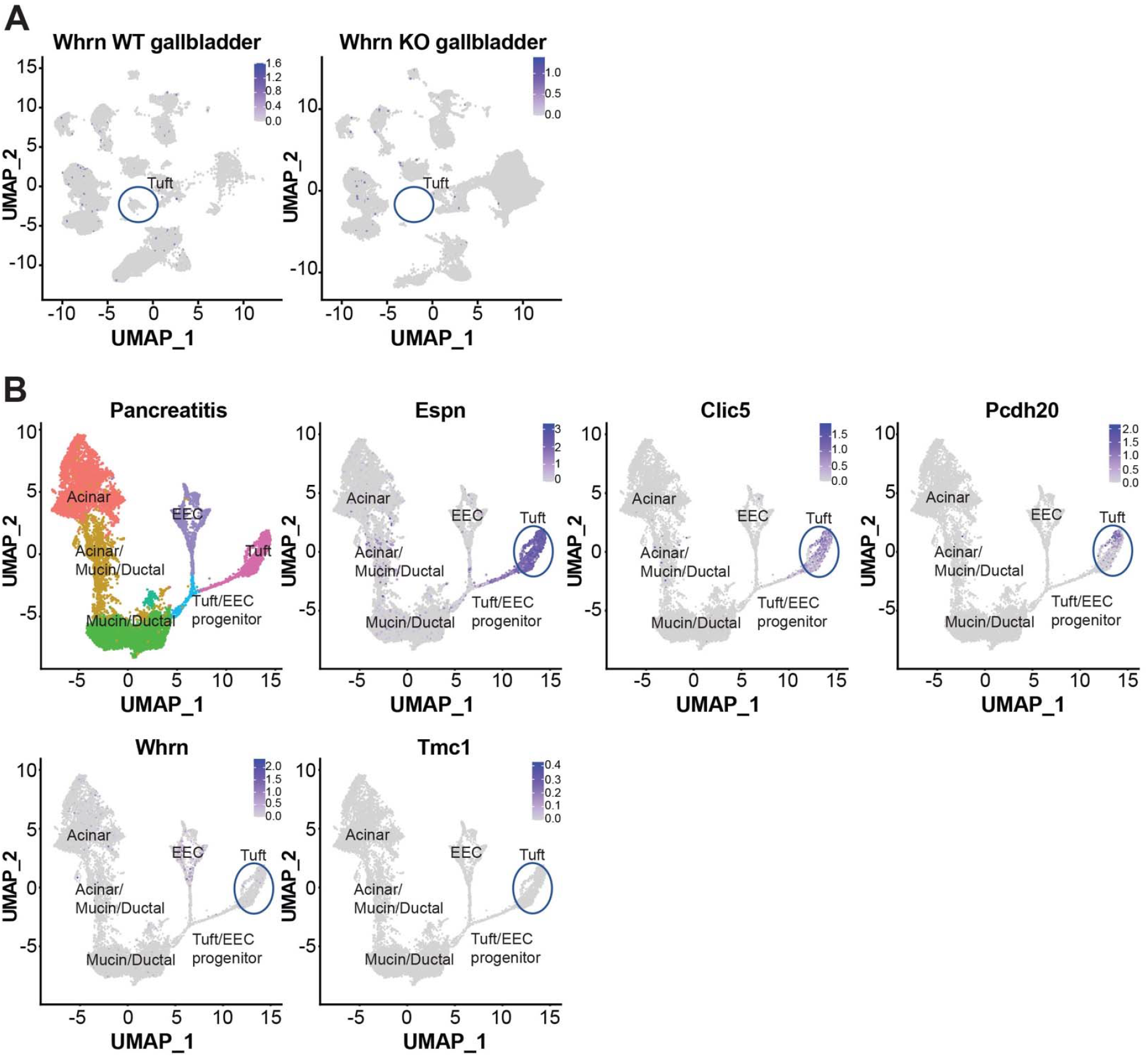
Expression of cochlear hair cell genes in gastrointestinal RNA-seq datasets. (**A**) Expression of *Whrn* in the gallbladders of either control or *Pou2f3*-/- mice generated from a study by O’Leary et al. (**B**) UMAPs of scRNA-seq data generated from murine pancreatitis from a study by Ma et al. Expression of *Espn, Clic5, Pcdh20, Whrn*, and *Tmc1* is highlighted in tuft cells.

**Figure S5.**
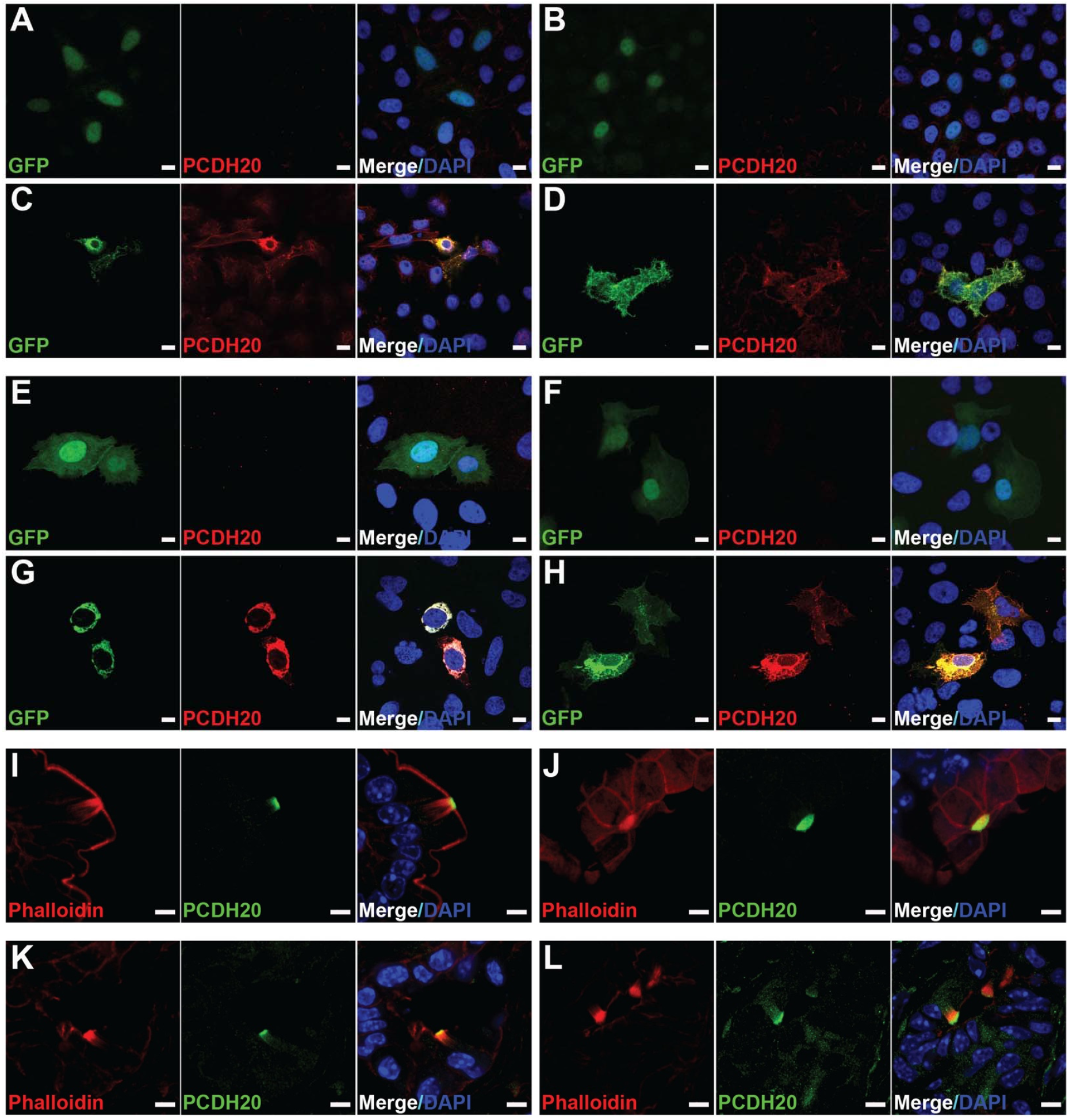
Validation of PCDH20 expression and localization. Transfection of either HeLA (**A-D**) or U2OS (**E-H**) cells with either GFP (green)(**A-B, E-F**) or PCDH20-GFP (green)(**C-D, G-H**), co-stained with an antibody against PCDH20 (red). DAPI, blue. Scale bars, 10 μm. IF for PCDH20 (green), phalloidin (red), and DAPI (blue) showing apical expression of PCDH20 in the microvilli of tuft cells from either the (**I-J**) small intestines or (**K-L**) pancreatic neoplasia. Scale bars, 5 μm. *In vivo* staining is representative of a minimum of n = 3 mice.

**Figure S6.**
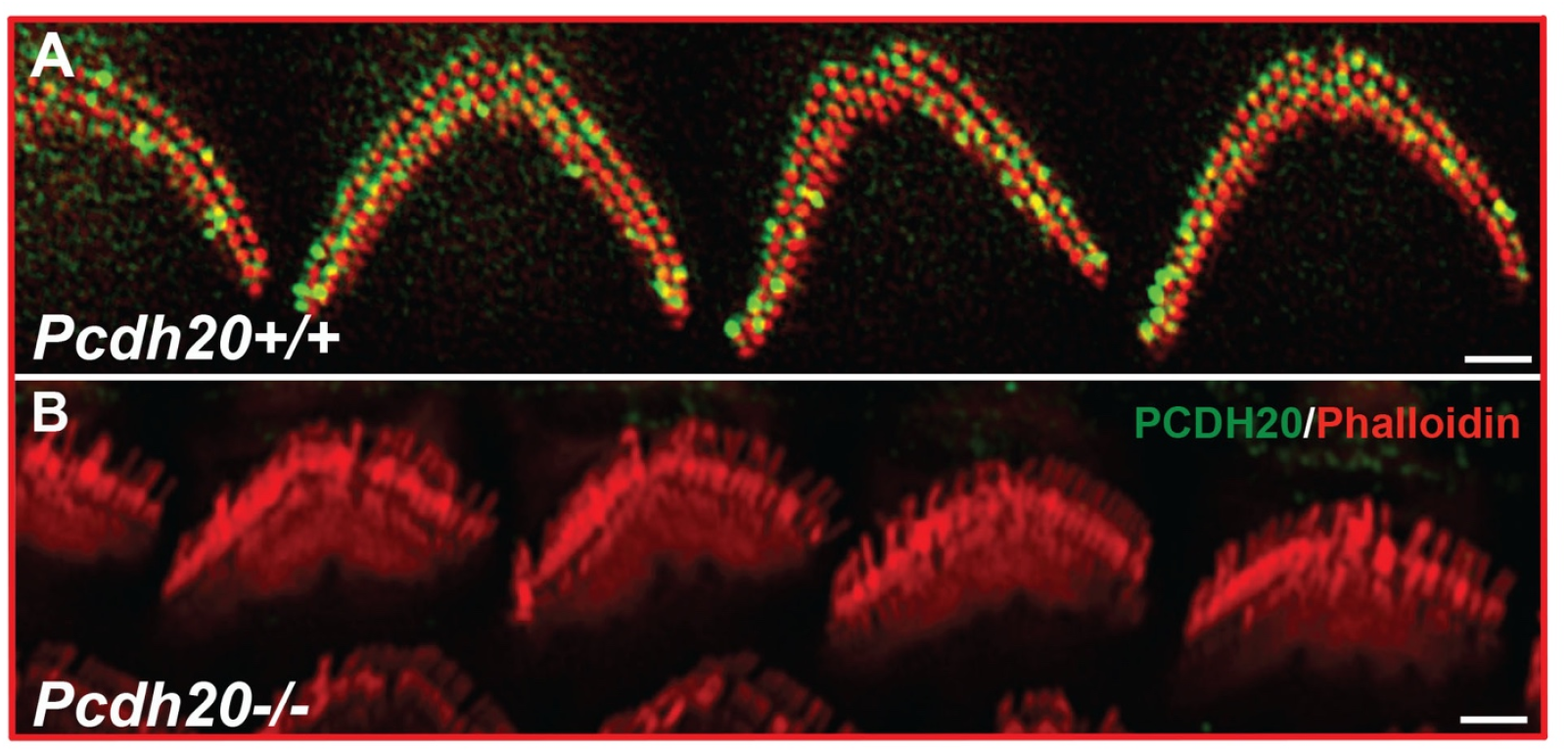
PCDH20 is expressed in hair cell stereocilia. Immunofluorescence for PCDH20 (green) with F-actin stained with phalloidin (red) in the cochlear inner hair cells of a (**A**) *PCDH20+/+* adult mouse indicating an inter-stereociliary localization, and (**B**) a *PCDH20-/-* adult mouse with no labeling. Scale bars = 1μm. n = 1 mouse/condition.

**Figure S7.**
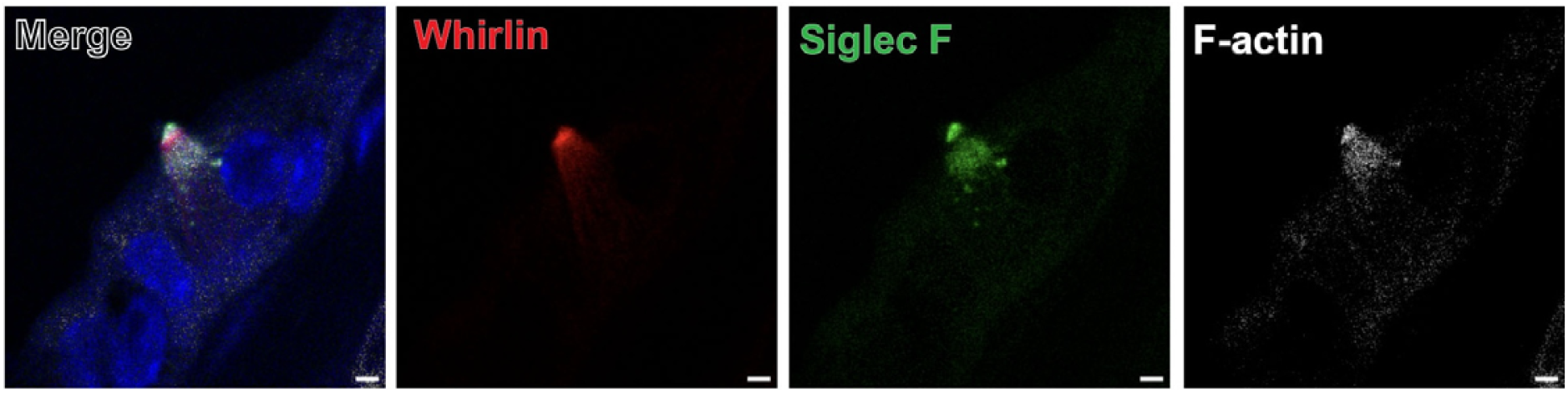
Whirlin expression in tuft cells. Co-immunofluorescence for whirlin (red), Siglec f (green), and F-actin (phalloidin, white) in a gallbladder tuft cell. DAPI, blue. Scale bars, 2 μm. Representative images are reflective of a minimum of n = 3 mice analyzed.

**Figure S8.**
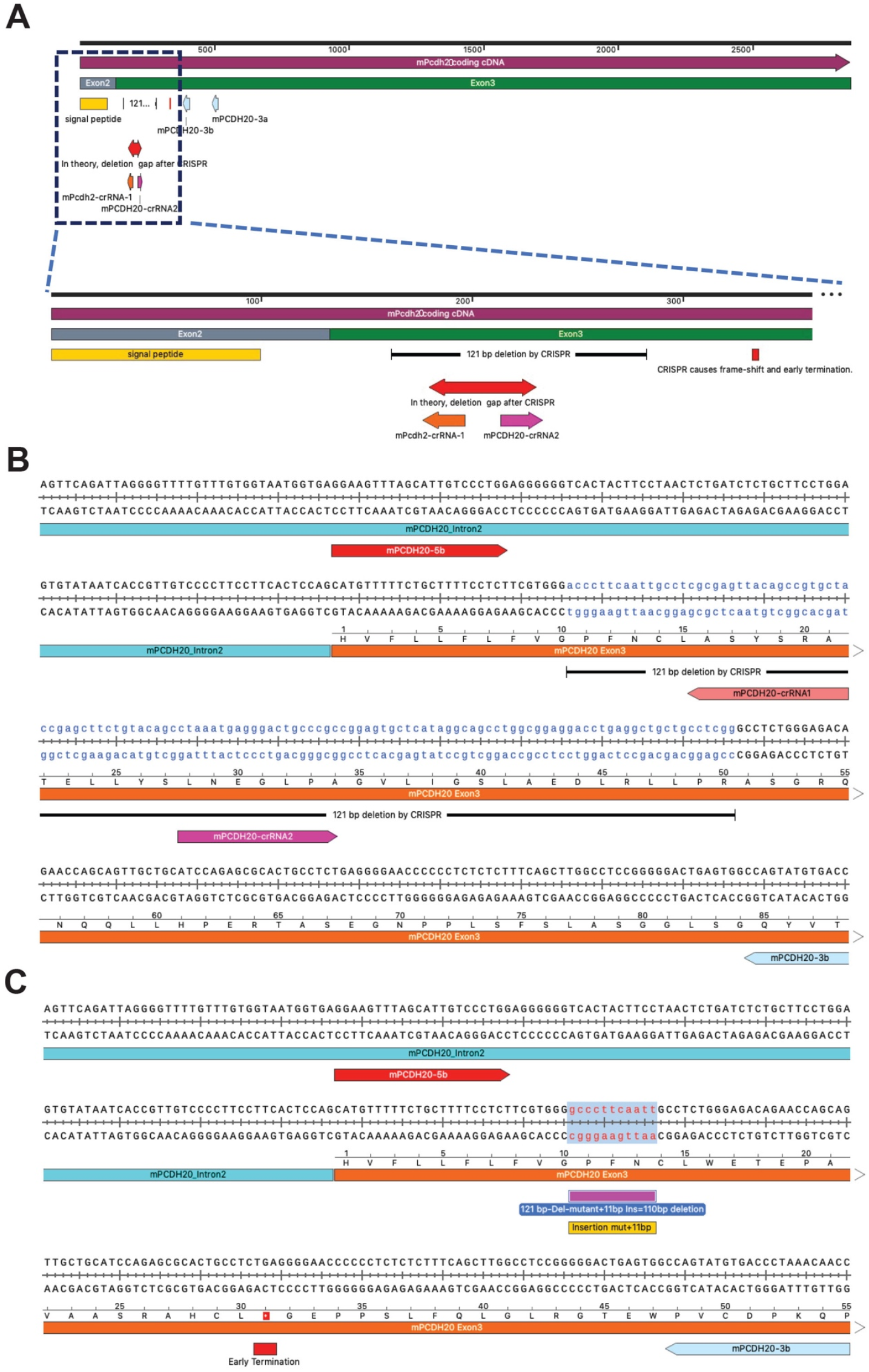
Generation of PCDH20 knockdown mice. (**A**) CRISPR strategy to generate PCDH20 knockdown mice. (**B**) DNA sequence of the wild-type *Pcdh20* gene with regions modified by CRISPR highlighted. (**C**) *Pcdh20* gene DNA sequence post-CRISPR modification highlighting a 121 base pair deletion and 11 base pair insertion.

**Figure S9.**
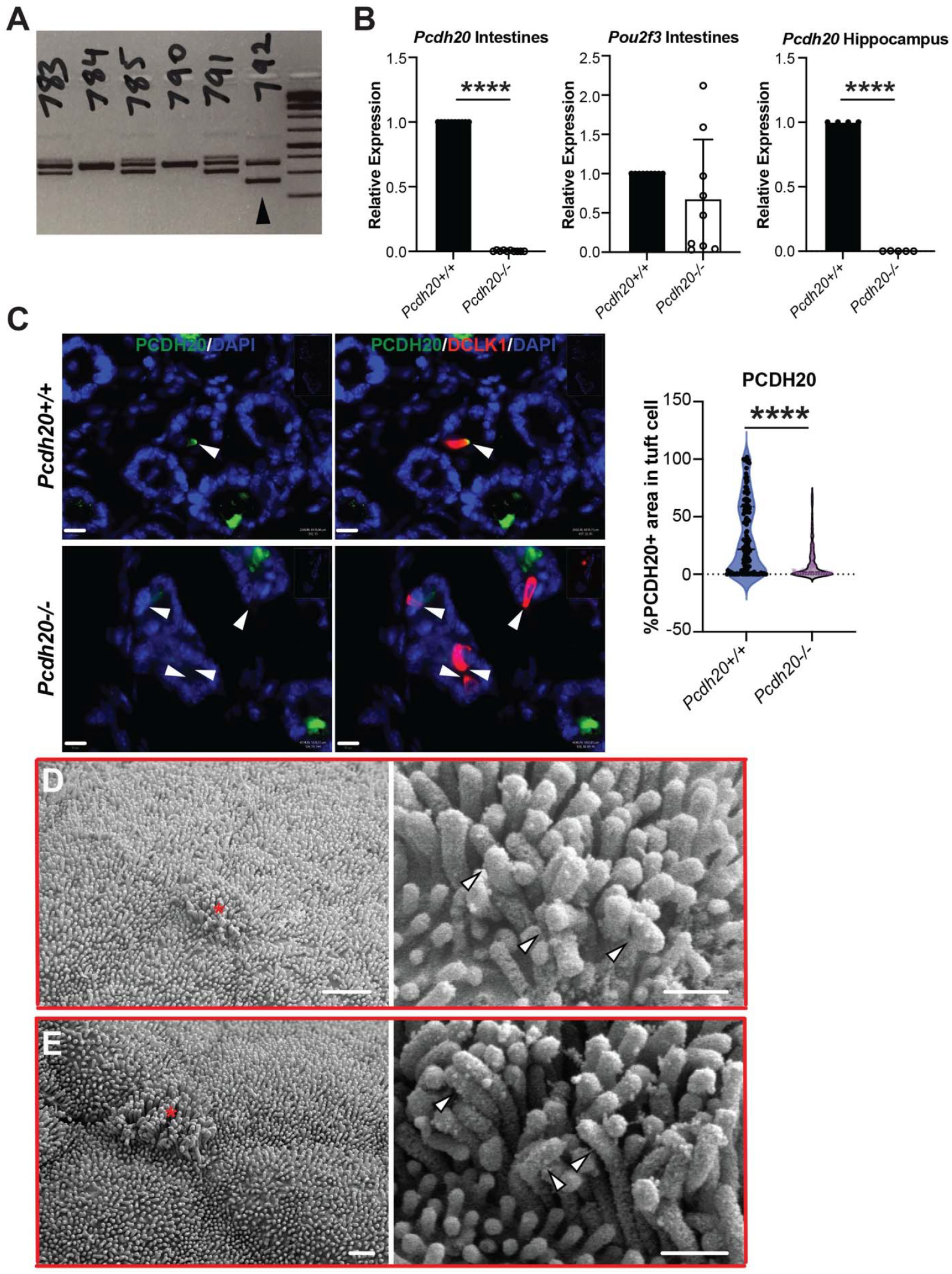
Analysis of PCDH20 knockdown mice. (**A**) Gel electrophoresis image of genotyping of multiple CRISPR mice screened for *Pcdh20* loss. Arrow, mouse line 792 selected for further analysis. (**B**) qRT-PCR analysis of small intestines or hippocampus for either the excised region of *Pcdh20* or *Pou2f3*. N = 4-6 mice. (**C**) Co-immunofluorescence and quantification of PCDH20 (green) as a % area of DCLK1+ (red) tuft cells in the small intestines of either *Pcdh20+/+* or *Pcdh20-/-* mice. n = 3 mice/genotype. DAPI, blue. Scale bars, 10 μm. Arrows, tuft cell microvilli. ^****^, p < 0.001. (**D-E**) SEM of gallbladder tuft cells from a *Pcdh20*-/- mouse. Scale bar, 1 μm, insert, 0.5 μm. Red asterisk, tuft cell. Arrows, bent microvilli.

**Figure S10.**
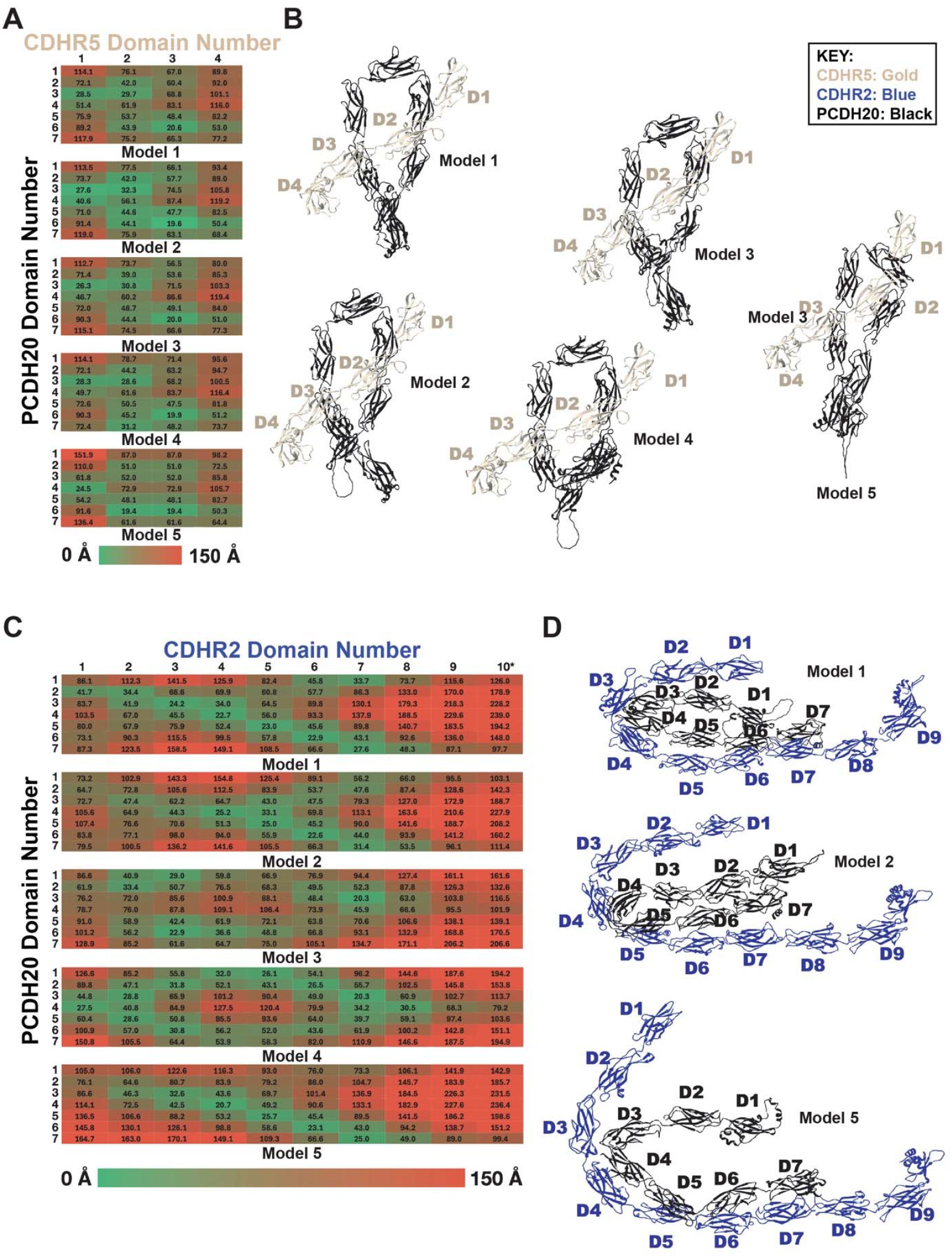
Potential Cadherin-PCDH20 interactions. (**A**) Heat map matrix displaying the center-to-center distance between each domain in CDHR5 (horizontal axis) and PCDH20 (vertical axis) for models 1 through 5. (**B**) Ribbon diagrams of interactions between cadherin domains 1-4 of CDHR5 (gold) with cadherin domains 1-7 of PCDH20 (black) are displayed for each model. Only non-sensical orthogonal interactions are present with parallel and antiparallel domain-domain interactions absent. (**C**) Heat map matrix displaying the center-to-center distance between each domain in CDHR2 (horizontal axis) and PCDH20 (vertical axis) for models 1 through 5. (**D**) Ribbon diagrams of interactions between cadherin domains 1-9 of CDHR2 (blue) with cadherin domains 1-7 of PCDH20 (black) are displayed for models 1, 2, and 5. Heat maps are colored from green (0 Å) to red (150 Å).

**Figure S11.**
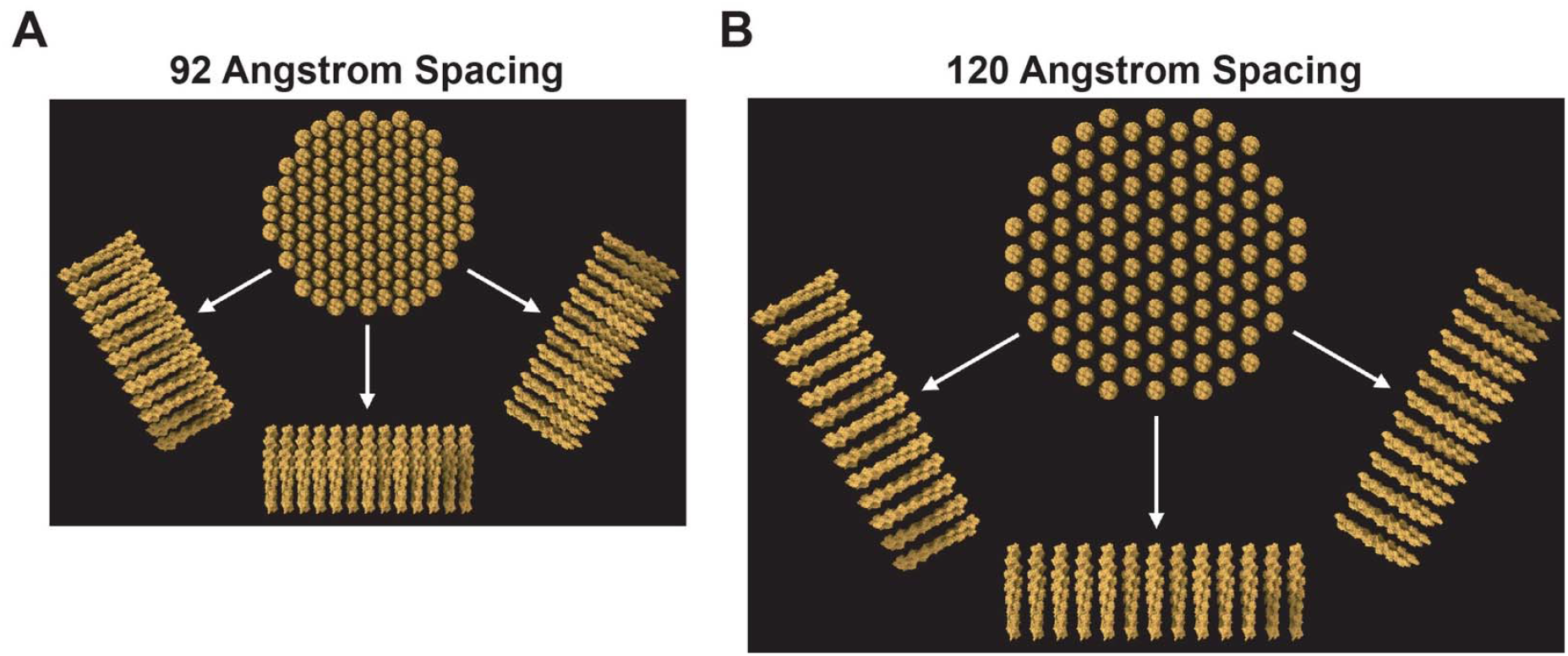
Actin bundles with different center-to-center spacing. (**A**) Molecular model of 109 hexagonally arranged actin filaments with a center-to-center spacing of 92 Å consistent with that measured by electron microscopy for the tuft cell cytoskeleton. The resulting bundle has a diameter of ∼100 nm, consistent with that measured for the tuft cell microvillus. (**B**) Molecular model of 109 hexagonally arranged actin filaments with a center-to-center spacing 120 Å, resulting in a bundle that has a diameter of 130 nm, far larger than the ∼106 nm median size. The actin filaments are modeled here as a fully saturated hexagonal lattice. Experimental data suggests that filaments are absent and that, on average, each actin filament has 5 (not 6) hexagonally arranged neighbors.

**Supplemental Video 1**. Clipping series through the actin bundle with center-to-center spacing 92 Å demonstrates there are no steric interactions between adjacent filaments.

